# Loss of SARM1 protects against retinal ganglion cell degeneration in Autosomal Dominant Optic Atrophy

**DOI:** 10.1101/2024.09.30.615936

**Authors:** Chen Ding, Papa S. Ndiaye, Sydney R. Campbell, Michelle Y. Fry, Jincheng Gong, Sophia R. Wienbar, Whitney Gibbs, Philippe Morquette, Luke H. Chao, Michael Tri H. Do, Thomas L. Schwarz

**Affiliations:** F.M. Kirby Neurobiology Center, Boston Children’s Hospital, Boston, MA 02115, USA; Department of Neurobiology, Harvard Medical School, Boston, MA 02115, USA; Department of Molecular Biology, Massachusetts General Hospital, Boston, MA 02114, USA; Department of Genetics, Blavatnik Institute, Harvard Medical School, Boston, MA 02115, USA; Department of Neurology, Harvard Medical School, Boston, MA 02115, USA

**Keywords:** Autosomal Dominant Optic Atrophy (ADOA), Retinal ganglion cell (RGC) degeneration, Mitochondria, OPA1, SARM1

## Abstract

Autosomal Dominant Optic Atrophy (ADOA), the most prevalent inherited optic neuropathy, leads to retinal ganglion cell (RGC) degeneration and vision loss. ADOA is primarily caused by mutations in the *OPA1* gene, which encodes a conserved GTPase important for mitochondrial inner membrane dynamics. To date, the disease mechanism remains unclear, and no therapies are available. Here, we present a novel mouse model carrying the pathogenic *Opa1^R290Q/+^* allele that recapitulates key features of human ADOA, including mitochondrial defects, age-related RGC loss, optic nerve degeneration, and reduced RGC functions. We identify SARM1, a neurodegeneration switch, as a key driver of RGC degeneration in these mice. *Sarm1* knockout nearly completely suppresses all the degeneration phenotypes. Additionally, we show that SARM1 is located within the mitochondrial intermembrane space (IMS). These findings indicate that SARM1 is activated downstream of mitochondrial dysfunction in ADOA, highlighting it as a promising therapeutic target.

## Introduction

Autosomal Dominant Optic Atrophy (ADOA) is one of the most common inherited optic neuropathies, affecting approximately 1 in 30,000 individuals worldwide.^1,2^ ADOA primarily causes degeneration of retinal ganglion cells (RGCs) and their axons, which form the optic nerve.^3^ Vision loss in ADOA patients typically begins insidiously during the first decade of life, with symptoms that usually progress slowly and can lead to legal blindness in severe cases.^4^ At least 70% of ADOA cases are caused by mutations in *OPA1,*^2,5–8^ a nuclear gene encoding a highly conserved GTPase essential for inner mitochondrial membrane fusion and cristae maintenance.^9,10^ Homozygous loss of *OPA1* is embryonic lethal, and most ADOA patients are heterozygous carriers exhibiting haploinsufficiency.^11^ However, certain missense mutations in the GTPase domain can act in a dominant negative fashion,^12,13^ and are often associated with a higher risk of developing a more severe syndromic disorder known as ADOA plus and characterized by additional symptoms such as deafness, ataxia, myopathy, peripheral neuropathy and progressive external ophthalmoplegia.^12,13^ Despite our understanding of the genetic underpinnings and efforts to build mouse models of ADOA,^14–16^ there are currently no treatments available, and our comprehension of its pathological mechanisms remains limited.

*OPA1* is tightly regulated at both the RNA and protein levels. Alternative splicing of exons 4b and 5b produces eight *OPA1* isoforms in humans, four in mice. The full-length OPA1 protein, which contains an N-terminal transmembrane domain, is embedded in the inner mitochondrial membrane. It undergoes proteolytic processing, with cleavage at the S1 site by the OMA1 protease^17,18^ and at the S2 and S3 sites by the YME1L protease,^19–21^ producing shorter forms that are released into the mitochondrial inner membrane space (IMS). Regulation of these protease activities^22,23^ maintains a balance between long and short OPA1 forms that cooperate to mediate membrane fusion and preserve cristae structure.^24–27^ Disruption of OPA1 function can lead to severe consequences, including fragmented mitochondria, abnormal cristae structure, reduced ATP production, increased oxidative stress, and loss of mitochondrial DNA (mtDNA).^28,29^

*SARM1*, a major pro-degenerative factor in neurons, encodes a conserved NADase that cleaves NAD^+^ into nicotinamide and cyclic ADPR.^30^ SARM1 has a mitochondrial targeting sequence, an N-terminal ARM domain followed by two SAM domains, and a C-terminal TIR domain. The TIR domain possesses intrinsic NAD^+^ cleavage activity but in healthy neurons is autoinhibited by the ARM domains.^31^ Under degenerative conditions, the autoinhibition is released by an elevated NMN/NAD^+^ ratio and triggers a positive feedback loop that rapidly consumes cytosolic NAD^+^.^32^ NAD^+^ depletion initiates a cascade of cellular events including ATP loss, calcium influx, and calpain activation that culminates in axon degeneration and neuronal cell death.^33^ *Sarm1* knockout (KO) can prevent degeneration in several neurodegenerative conditions, including glaucoma, traumatic brain injury, and peripheral neuropathy.^34–37^ Of particular interest is the role of SARM1 in degeneration induced by mitochondrial dysfunction, such as in Charcot-Marie-Tooth disease type 2A (CMT2A), which is caused by *mitofusin* mutations that disrupt mitochondrial outer membrane fusion.^38–40^ The connection between SARM1, mitochondrial dysfunction, and retinal degeneration prompted us to investigate whether SARM1 drives RGC degeneration in ADOA.

In this study, we developed and characterized a novel mouse model of ADOA by introducing the pathogenic *Opa1^R290Q/+^* mutation that has been reported in several ADOA families,^11,41,42^ and studied the impact of *Sarm1* deletion on the progression of optic atrophy in this mouse model.

## Results

### Generation of the *Opa1^R290Q/+^* mouse and allele characterization

We used the CRIPSR/Cas9 technique to introduce R290Q, a pathogenic mutation identified in human patients,^11,41,42^ into the *Opa1* locus of C57BL/6J mice (Figure S1A). Homozygous mice are embryonically lethal but heterozygotes are viable and fertile. The R290Q mutation is located in the first β-strand of the dynamin-type GTPase domain and included in all *Opa1* isoforms and cleavage products (Figures 1A and S1B). This mutation is distal from the nucleotide binding site and likely alters the folding of the GTPase domain core while leaving the C-terminal stalk and paddle regions intact.^43^ Unlike heterozygous alleles with early-stop codons that reduce OPA1 protein levels by half, the R290Q mutation appeared to alter OPA1 protein processing: lower molecular-weight isoforms were reduced and two higher molecular-weight bands appeared (Figures S1B-S1C). Similar changes were observed in human *OPA1^R290Q/+^* fibroblasts.^42^ The new bands likely correspond to full-length isoforms 5 and 8, which are completely cleaved into short forms and thus absent in WT controls.^19,21^ PCR analysis of *Opa1* isoforms in reverse-transcribed cDNAs from cultured *Opa1^R290Q/+^* fibroblasts and primary cortical neurons showed no detectable changes in *Opa1* transcript abundance (Figure S1D). The *OPA1^R290Q/+^* allele has dominant negative properties that can cause the ADOA plus phenotype in some human patients^41,42^. This may be due to the mutant protein integrating into potential OPA1 lattices together with the WT protein on the membrane through its stalk and paddle region,^44,45^ but exhibiting less GTPase-stimulated activity, leaving the fission forces unopposed.

**Figure 1.**
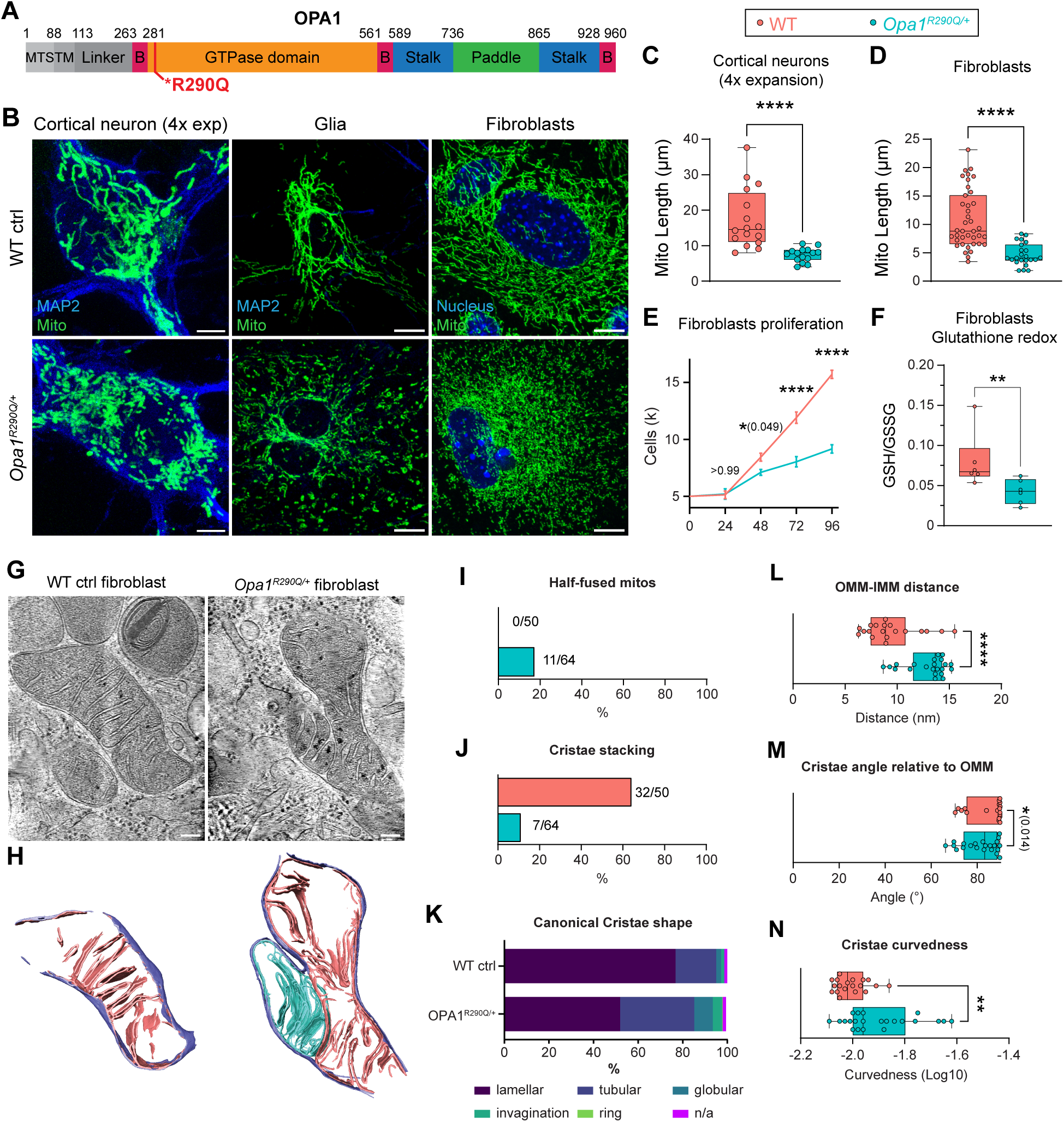
The *Opa1^R290Q/+^* mutation impairs mitochondrial morphology and cristae structure (A) Domain organization of mouse OPA1 isoform 1 based on the structure of human OPA1 isoform 1.^44^ The R290 residue is located at the beginning of the GTPase domain. MTS, Mitochondrial Targeting Sequence; TM, Transmembrane Domain; B, Bundle Signaling Element. (B) Representative confocal images of mitochondria in cortical neurons (DIV9), glial cells and fibroblasts isolated from *Opa1*^R290Q/+^ mice and WT littermate controls. In neurons and glia, mitochondria were labeled via lentiviral transduction of a mitochondrial matrix marker (COX8 pre-sequence fused to DsRed, or MitoDsRed). Expansion microscopy was performed on neurons. In fibroblasts, mitochondria were labeled by staining for endogenous ATP5α. Scale bars = 10 µm. (C-D) Quantification of mitochondrial length shows fragmented mitochondria in *Opa1^R290Q/+^* cortical neurons (C, n = 16 WT neurons and 15 *Opa1^R290Q/+^* neurons from 2 cultures) and fibroblasts (D, n = 39 WT cells from 2 experiments and 23 *Opa1^R290Q/+^* cells from 3 experiments). Mann-Whitney test. (E) *Opa1^R290Q/+^* fibroblasts (n = 6 wells) grow at a slower rate compared to the WT control (n = 6 wells). Mean ± SEM. Two-way ANOVA followed by Sidak multiple comparisons; effect of genotype (*F1, 50* = 109.7, *P* < 0.0001); time (*F4, 50* = 163.6, *P* < 0.0001); genotype x time (*F4, 50* = 32.44, *P* < 0.0001). (F) GSH/GSSG ratios in fibroblasts as measured by LC-MS/MS. N = 6 samples from 2 experiments for both genotypes. Mann-Whitney test. (G-H) Representative summed projections of the central slices of cryo-electron tomograms of WT and *Opa1^R290Q/+^* mitochondria (G) and corresponding 3D segmentations of the entire stack (H). In the *Opa1^R290Q/+^* cell, the OMM (purple in H) has fused while the IMMs (blue and pink in H) remain separate. Scale bars = 100 nm. (I) Percentage of mitochondria with fused OMM but separate IMMs. Numbers of stalled- fusion/total mitochondria are indicated. (J) Percentage of WT mitochondria (n = 50) and *Opa1^R290Q/+^* mitochondria (n = 64) displaying stacked cristae. (K) Classification of cristae morphology (n = 362 WT and 337 *Opa1^R290Q/+^* cristae). (L-N) The highest frequency values for OMM-IMM distance (L), cristae angle relative to the OMM (M), and cristae curvedness (N) for WT (n = 19) and *Opa1^R290Q/+^* (n = 22) mitochondria. Mann-Whitney test. Box plots denote minimum, first quartile, median, third quartile, and maximum values. \**P* < 0.05; \*\**P* < 0.01; \*\*\*\**P* < 0.0001.

### The *Opa1^R290Q/+^* mutation causes mitochondrial fragmentation, impairs mitochondrial function, and exhibits a dominant-negative effect

We first examined the impact of the *Opa1^R290Q/+^* mutation on mitochondrial morphology in primary cells isolated from the *Opa1^R290Q/+^* mice and their WT littermate controls. In cultured DIV9 cortical neurons, mitochondria were labeled via lentiviral transduction of MitoDsRed, and the network in the somata was resolved using a hydrogel-based expansion technique^46^ (Figure 1B). The mitochondrial network was fragmented in *Opa1^R290Q/+^* neurons, with average mitochondrial length reduced by 52% compared to WT (Figures 1B-1C). Similar fragmentation occurred in MAP2-negative glial cells in the cultures. In cultures of embryonic fibroblasts stained for endogenous ATP5α (Figures 1B and 1D), *Opa1^R290Q/+^* mitochondrial length was reduced to 43% of WT length. These phenotypes are consistent with the R290Q mutation severely impairing the fusion activity of OPA1, as previous reported in human fibroblasts.^42^

To determine whether the R290Q mutation is dominant negative, we overexpressed either WT OPA1 isoform 1 or OPA1 R290Q isoform 1 as a GFP fusion protein in WT fibroblasts (Figure S2A). While WT OPA1 overexpression did not significantly alter mitochondrial length, the OPA1 R290Q transgene reduced mitochondrial length by 40% (Figure S2B). Thus, while OPA1 overexpression *per se* has little effect, the R290Q allele can exert a dominant-negative effect, likely resulting in a more severe ADOA phenotype than a simple loss-of-function mutation.

In addition to morphological changes, the R290Q mutation also induced cellular changes consistent with mitochondrial dysfunction. *Opa1^R290Q/+^* fibroblasts proliferated at a slower rate compared to WT controls (Figure 1E) and had a significantly lower GSH/GSSG (reduced/oxidized) ratio as measured by LC-MS/MS,^47^ indicating cellular oxidative stress (Figure 1F).^48^ The same change in glutathione redox was observed in whole brain metabolites (Figure S3A). In the Seahorse assay (Agilent) of mitochondrial respiration, mutant fibroblasts showed slightly reduced baseline oxygen consumption rates and proton leakage, with minimal changes in maximum respiration (Figures S3C-S3D). Additionally, no changes were detected in mtDNA copy numbers, as measured by RT-qPCR using three mitochondrial genes (Figure S3B).

### Mitochondria in *Opa1^R290Q/+^* fibroblasts have aberrant cristae

In addition to mediating membrane fusion, OPA1 plays an important role in shaping mitochondrial cristae.^10^ We performed cryo-electron tomography (Cryo-ET) on cryo-focused ion beam (cryo-FIB)-milled fibroblasts to examine their 3D cristae ultrastructure. We focused on the fibroblasts due to their homogeneity and abundant mitochondria. WT and *Opa1^R290Q/+^* fibroblasts were deposited on grids, back-blotted, and vitrified in liquid ethane. Windows in the cells were milled with a cryo-focused ion beam scanning electron microscope (cryo-FIB-SEM) to generate 150-200 nm thick lamellae, where mitochondria were targeted and tilt-series were collected in a 300 keV Titan Krios and processed to generate 3D tomograms (Figures 1G-1H).

When examining gross mitochondrial morphology, we observed a notable fusion defect: 11 out of 64 *Opa1^R290Q/+^* mitochondria exhibited a fused outer membrane (OMM) but separated inner membrane (IMM). This half-fused state was not observed in WT mitochondria (Figures 1H-1I), and this defect, specifically lacking inner-membrane fusion, is consistent with impaired OPA1 function. We interpret these mitochondria as having stalled IMM fusion, rather than being in the process of fission, based on two observations: a) the absence of endoplasmic reticulum (ER), actin or septin filaments proximal to the contact site of the two mitochondria;^49,50^ and b) the lack of any potential DRP1 densities corresponding to a constriction ring at the OMM.^51,52^ Furthermore, IMM severing likely occurs either simultaneously or shortly after OMM severing during fission, making visualizing intermediates unlikely. We propose the observation of two distinct IMMs with a fused OMM in the *Opa1^R290Q/+^* mitochondria represents an intermediate state in the fusion process, since fusion of the outer and inner membrane occurs sequentially and has been previously shown to be decouplable in cells and *in vitro*.^53,54^

We also noted differences in cristae architecture in the *Opa1^R290Q/+^* mitochondria. A reduction in three or more parallel stacked cristae was observed in *Opa1^R290Q/+^* mitochondria (Figure 1J).

Cristae were classified into the following canonical shape categories: lamellar (sheet-like), tubular, globular (balloon-like), ring, invagination (short projections into the mitochondria matrix), or undetermined (Figure 1K). A reduction in lamellar cristae and increase in tubular cristae were observed in *Opa1^R290Q/+^* mitochondria, consistent with the observed decrease in stacked cristae.

Cristae architectural differences were further quantified by applying a morphometrics toolkit to analyze segmented membranes from mitochondria in WT (n = 19 mitochondria) and *Opa1^R290Q/+^* (n = 22 mitochondria) cells.^55^ One notable change was that the distance between the OMM and the IMM was larger in *Opa1^R290Q/+^* mitochondria compared to wild type (Figure 1L). This suggests that, in addition to impairing fusion, the mutation disrupts the machinery that maintains the spacing between inner and outer membranes. In mutant mitochondria, cristae were less perpendicular relative to the OMM and more curved compared to WT mitochondria (Figures 1M and 1N). Thus, these Cryo-ET data reveal that the *Opa1^R290Q/+^* mutation impairs cristae ultrastructure in addition to its adverse effects on mitochondrial networks and morphology.

### Age-related retinal ganglion cell (RGC) death and optic nerve degeneration in *Opa1^R290Q/+^* mice

Retinal ganglion cells (RGCs) are the primary cells affected by ADOA. To study RGC death and survival, we examined retinal whole mounts from a large cohort of *Opa1^R290Q/+^* mice and their littermate WT controls, with equal representation of the sexes, and ranging from 3 to 18 months of age at 3-month intervals. Retinas were stained for RBPMS, a pan-RGC marker,^56^ and phospho-H2Ax, a histone marker for DNA double-strand breaks, which labels RGCs undergoing cell death^57^ (Figure 2A, arrowheads highlight phospho-H2Ax-positive RGCs). In 3- and 6-month-old (MO) animals, WT and *Opa1^R290Q/+^* retinas had similar numbers of RGCs and percentages of dying RGCs (Figures 2A-2C). However, by 9 months of age, *Opa1^R290Q/+^* mice displayed a 3-fold increase in dying RGCs compared to age-matched WT controls, although this was not yet reflected in a significant decrease in the total RGC count (Figures 2A-2C). At 12 months, a reduction in total RGCs was apparent in *Opa1^R290Q/+^* mice alongside an increase in dying RGCs. By 15 months, the mean total RGC number had decreased by 16% compared to WT, with dying RGCs rising to 4-fold of the WT level (Figures 2A-2C). At 18 months, the difference in dying RGCs was largely diminished, and there was no further decrease in the total RGC number (Figures 2B-2C). Taken together, these findings indicate that retinas develop normally in *Opa1^R290Q/+^* mice and that subsequent RGC death predominantly occurs between 9 and 15 months of age. The progressive death of RGCs is consistent with the slow progressive nature of vision decline in human patients^58^.

**Figure 2.**
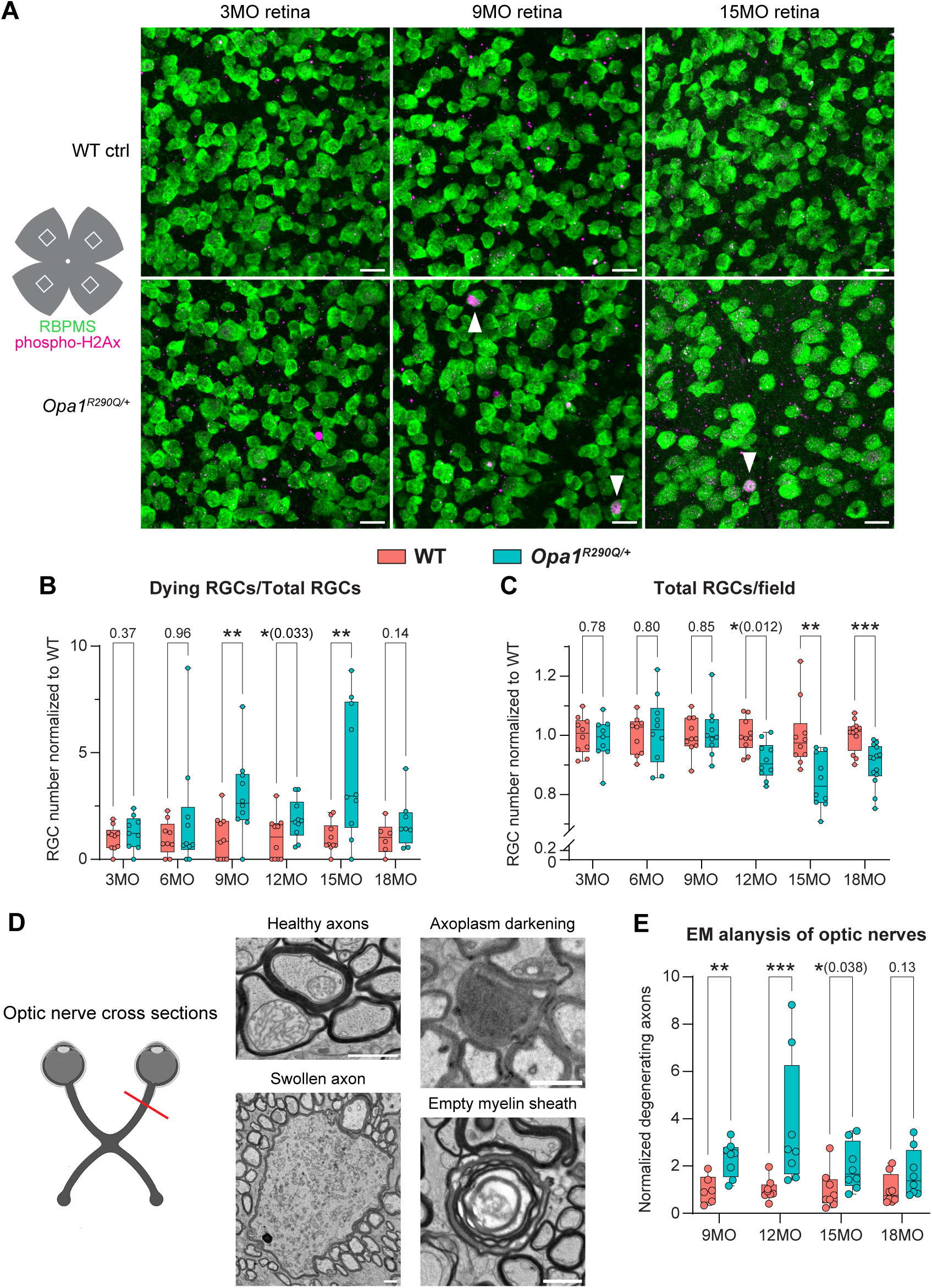
Age-related RGC and optic nerve degeneration in *Opa1^R290Q/+^* mice (A) Representative images of retinal whole mounts from WT and *Opa1^R290Q/+^* mice. Each retina was imaged in every quadrant, approximately 1.2 mm from the center. RBPMS (green) marks all RGCs and phospho-H2Ax (magenta) labels RGCs undergoing cell death (marked by arrowheads). Scale bars = 20 µm. (B) Quantification of Dying RGCs over time. Each dot represents one retina. The counts of dying RGCs were averaged across the four quadrants of each retina and divided by the average RGC number of those quadrants. This value was then normalized to the WT average at each age. N = 9-12 WT and 9-14 *Opa1^R290Q/+^* retinas per age group. (C) Quantification of total RGCs over time. Each dot represents one retina. RGC counts were first averaged across the four quadrants of each retina and then normalized to the WT average at each age. N = 6-10 WT and 8-10 *Opa1^R290Q/+^* retinas per age group. (D) Diagram of optic nerve cross-sections and representative EM images. Healthy axons have compact myelin sheaths and microtubules in the axoplasm. Degenerating axons were classified into three categories: swollen axons with accumulation of organelles or neurofilaments, dark axons in the late stage of degeneration, and empty myelin sheath with completely degenerated axons. Scale bars = 1 µm. (E) Quantification of degenerating axons. Each dot represents one optic nerve. The counts of degenerating axons were averaged across the nine fields of each cross-section of an optic nerve and divided by the average axon number of those fields. This value was then normalized to the WT average at each age. N = 6-8 WT and 8 *Opa1^R290Q/+^* retinas per age group. Box plots denote minimum, first quartile, median, third quartile, and maximum values. \**P* < 0.05; \*\**P* < 0.01; \*\*\**P* < 0.001 determined by Mann-Whitney test for each age group.

Electron microscopy (EM) images of optic nerve cross-sections indicated progressive degeneration of RGC axons. Healthy axons are characterized by well-aligned microtubules, and a compact myelin sheath (Figure 2D). In *Opa1^R290Q/+^* mutants, we observed that a fraction of the axons were undergoing recognizable stages of degeneration: early-stage degeneration included axon swelling and accumulation of organelles or neurofilaments, followed by axoplasm darkening in the late stage, and eventually leading to fully degenerated axons with only empty myelin sheaths remaining (Figure 2D). Between 9 and 15 months, the fraction of degenerating axons was significantly higher in mutants compared to WT (Figure 2E). By 18 months, the difference was no longer significant, consistent with the observed pattern of RGC cell death (Figure 2B).

### Visual evoked potentials decline with age in *Opa1^R290Q/+^* mice

To assess whether RGC degeneration leads to vision deficits in *Opa1^R290Q/+^* mice, we performed longitudinal recordings of dark-adapted electroretinograms (ERG) and visual evoked potentials (VEP) in a large cohort of mice (16 per genotype, 8 males and 8 females). Two LED stimulators with built-in electrodes were placed in close contact with the eyes of anesthetized mice to deliver light stimuli and record ERGs from the cornea. A subcutaneous needle electrode was positioned along the midline above the visual cortex to capture VEPs, while a reference electrode was placed in the snout, and a ground electrode was inserted under the skin near the tail (Figure 3A). This setup allowed for non-invasive, simultaneous recordings of ERG and VEP in the same cohort of mice as they aged (example responses shown in Figure 3B).

**Figure 3.**
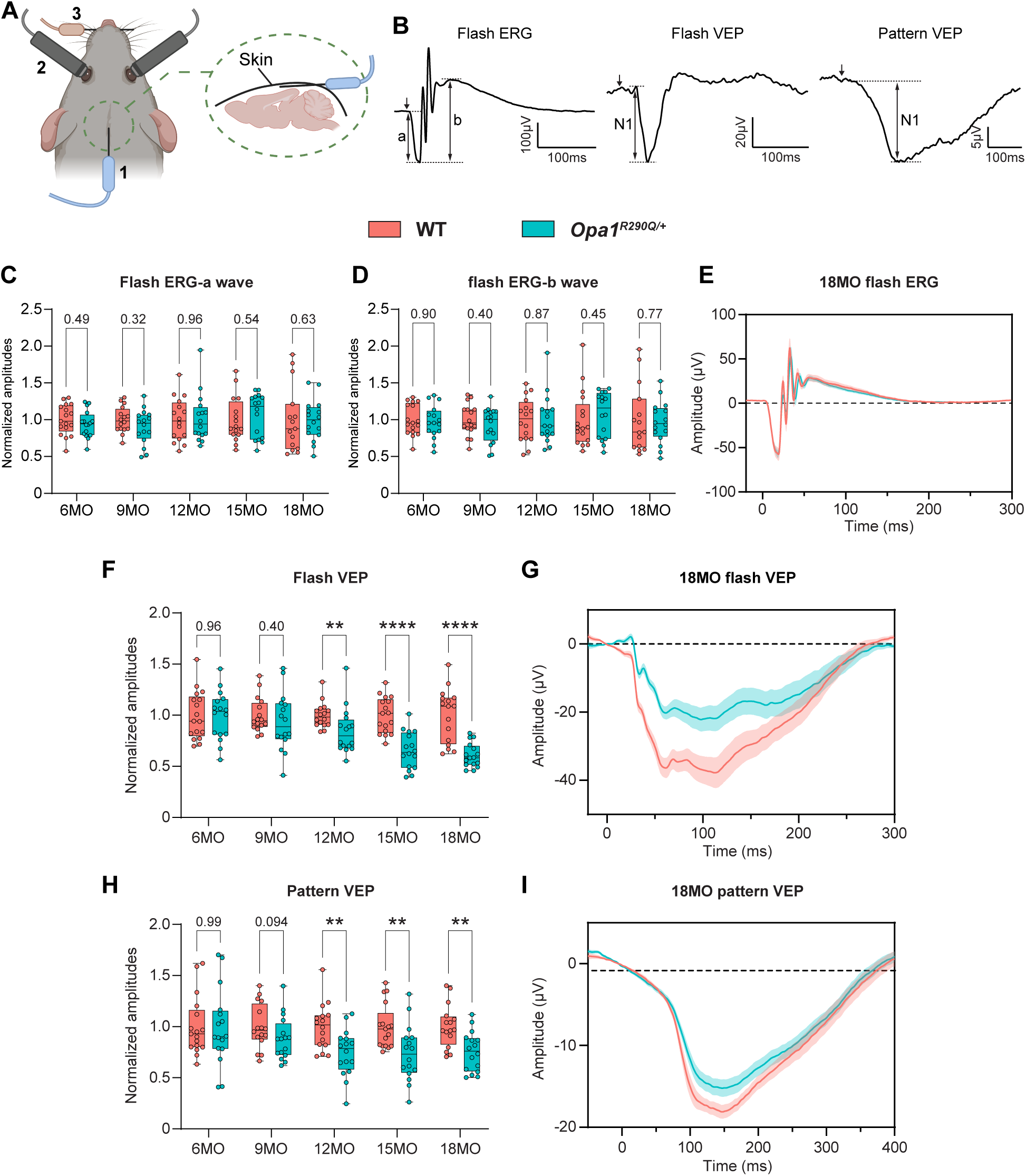
Impaired RGC function in *Opa1^R290Q/+^* mice as measured by electroretinogram and visual evoked potential (A) Diagram of recording setup. 1, subcutaneous recording electrode for VEP; 2, Integrated LED stimulator and recording electrode for ERG; 3, Snout reference electrode for VEP. A ground electrode was inserted subcutaneously next to the tail (not shown). (B) Representative ERG and VEP traces from WT control mice. For flash ERG, the amplitudes of a-wave and b-wave were measured; For flash and pattern VEPs, the amplitude of N1 was measured. (C-D) Amplitudes of a-wave (C) and b-wave (D) of flash ERG across age. Each dot represents one animal. Amplitudes were normalized to WT average at each age. N = 15-17 WT and 16 *Opa1^R290Q/+^* mice per age group. (E) Average flash ERG traces in 18MO animals. N = 15 WT and 16 *Opa1^R290Q/+^* mice. Mean ± SEM. (F) N1 amplitudes of flash VEP across age. Each dot represents one animal. Amplitudes were normalized to WT average at each age. N = 16-17 WT and 16 *Opa1^R290Q/+^* mice per age group. (G) Average flash VEP traces in 18MO animals. N = 16 WT and 16 *Opa1^R290Q/+^* mice. Mean ± SEM. (H) N1 amplitudes of pattern VEP across age. Each dot represents one animal. Amplitudes were normalized to WT average at each age. N = 16-17 WT and 16 *Opa1^R290Q/+^* mice per age group. (I) Average pattern VEP traces in 18MO animals. N = 16 WT and 16 *Opa1^R290Q/+^* mice. Mean ± SEM. Box plots denote minimum, first quartile, median, third quartile, and maximum values. \*\**P* < 0.01; \*\*\*\**P* < 0.0001 determined by Mann-Whitney test for each age group.

For ERGs, light flashes evoke responses from a mixed neuronal population in the retina, excluding RGCs. The response waveform is characterized by a negative a-wave, followed by oscillations and a positive b-wave.^59,60^ The a-wave is generated by photoreceptors, while the b-wave, measured from trough to peak, is primarily produced by bipolar cells, with potential contributions from Müller glia^60,61^ (Figure 3B). The amplitudes of both the a-wave and b-wave of the flash ERG (fERG) did not differ between WT and *Opa1^R290Q/+^* mice at any of the ages examined (Figures 3C-3D). The average fERG traces in 18MO animals were nearly identical in the two genotypes (Figure 3E), consistent with the selective degeneration of RGCs in ADOA.

Visual evoked potentials originate from the visual cortex and depend on the ability of RGCs to transmit electrical signals to the cortex via the geniculo-cortical pathway. As such, VEPs are widely used to assess RGC connections to the brain.^62–64^ We measured VEP responses using flash stimuli (fVEP) and patterned stimuli (pVEP) consisting of alternating horizontal bars. Each type of stimulus produces a stereotypical waveform with a negative peak N1 (Figure 3B).

Quantification of the N1 amplitudes revealed no differences between WT and *Opa1^R290Q/+^* mice at or before 9 months of age, suggesting normal visual development and function in the *Opa1* mutants. However, starting at 12 months, the mutant mice showed decreased responses (Figures 3F and 3H). By 18 months, the N1 amplitudes of fVEPs were reduced to 61% of WT levels, and pVEP N1 amplitudes were reduced to 75% of WT levels. The average response traces also showed clear separation between 18MO WT and *Opa1^R290Q/+^* mice (Figures 3G and 3I). The timing of VEP decline closely aligns with the trajectory of RGC loss and optic nerve degeneration (Figures 2C and 2E). Collectively, these data suggest that RGCs in *Opa1^R290Q/+^* mice begin to degenerate detectably around 9 months of age, leading to vision defects starting at 12 months.

To examine whether RGC degeneration leads to behavioral deficits, we assessed visual function using the optomotor reflex (OMR) assay. Mice were placed in a computer-monitored arena surrounded by screens displaying moving sine wave gratings at varying spatial frequencies and contrasts, while reflexive head tracking of the gratings was automatically quantified (qMOR, PhenoSys). However, we did not detect significant changes in OMR responses to varying spatial frequencies or contrasts regardless of the age of the mice, even in 18MO *Opa1^R290Q/+^* mice (Figure S4). OMR is primarily mediated by ON direction-selective RGCs (ON DSGCs), which project to the accessory optic system (AOS)^65–67^ and may constitute only about 10% of the total RGC population in the mouse retina.^68^ Given that OMR in response to vertical gratings likely depends on an even smaller subset of ON DSGCs that respond to temporal-to-nasal motion, it is plausible that this subset is not part of the degenerating population of RGCs, or that too few of them are lost to alter the behavioral response.

### *Opa1^R290Q/+^* mice exhibit altered compound action potentials in the optic nerve

To examine RGC function in a more direct and controlled manner, we isolated the retina and attached optic nerve from 20-21MO animals, delivered light to the retina, and recorded signals from the cut end of the nerve using a suction electrode (Figure 4A). These signals are understood to reflect the summed action potentials of retinal ganglion cells (RGCs): the compound action potential (CAP) (Figure S5A).^69^ To our knowledge, light-evoked CAPs have not been reported for the *ex vivo* optic nerve of the mouse. To verify that CAPs originated from RGC action potentials, we conducted pharmacological experiments. CAPs were reversibly blocked by tetrodotoxin, an antagonist of voltage gated sodium channels and thus RGC action potentials (Figure S5B). CAPs were also blocked by antagonists of synaptic transmission, indicating a reliance on signaling from rod and cone photoreceptors (Figure S5B). The intrinsic photosensitivity of melanopsin-expressing RGCs^70^ was not evident, perhaps due to the low abundance of these neurons. These observations are consistent with CAPs comprising the population action potentials of RGCs driven by rods and cones.

**Figure 4.**
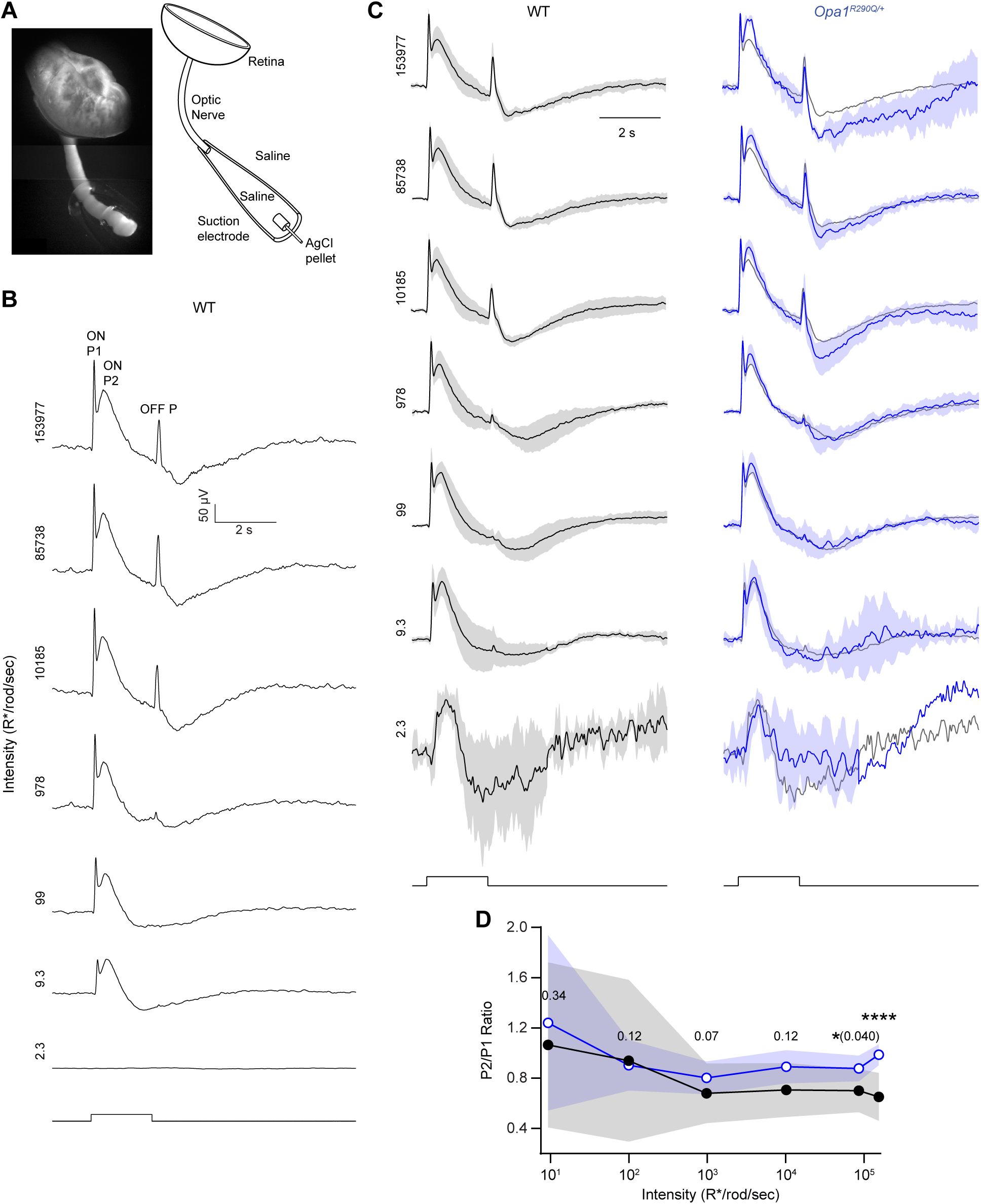
Abnormal compound action potentials (CAPs) in the optic nerve of *Opa1^R290Q/+^* mice (A) *Left*, The recording configuration contains the retina (top) and optic nerve (center), with the latter drawn into an electrode. *Right*, Schematic showing the retina, nerve and electrode. (B) Example CAPs from a WT retina. For lower intensities (<10 R*/rod/sec), at least 3 trials were typically averaged to increase signal/noise; for brighter intensities, one trial sufficed. Stimulus monitor is illustrated at the bottom, with light intensities increasing from bottom to top (indicated on the left in units of R*/rod/sec). (C) CAPs (Mean ± SD) of WT (left, black) and *Opa1^R290Q/+^* (right, blue) retinas. Responses of each retina were normalized to its maximum value at each light intensity. The normalized *Opa1^R290Q/+^* traces (blue) were superimposed on the mean of the normalized WT traces (gray) for comparison. Shorter trials were used for dimmer stimuli, leading to a lack of error bars in some intervals. N = 7 WT and 7 *Opa1^R290Q/+^* retinas per age group, except for 3 *Opa1^R290Q/+^* retinas at the highest intensity. (D) Ratio of second and first ON peaks (ON P2/P1) from (C). Mean ± SD shown for WT (black) and *Opa1^R290Q/+^* (blue). For the dimmest intensity, distinct peaks were not evident (average Z- score < 10) and were therefore not included in analyses. \**P* < 0.05; \*\*\*\**P* < 0.0001 determined by Mann-Whitney test with bootstrapping (see methods).

To evaluate the effect of *Opa1^R290Q/+^* on the generation and propagation of CAPs, we delivered pulses of light (2 s duration) that evoked activity during light onset (ON responses) and offset (OFF responses) (Figure 4B). We began at an intensity that is suprathreshold for rod-driven responses in many RGC types (1 R*/rod/s)^71^ or below. We increased the intensity until reaching saturation or near-saturation of the response (∼1.5×10^5^ R*/rod/s). We set the inter-pulse interval to avoid cumulative desensitization (i.e., responses at a given intensity showed no progressive change; <0.75% rhodopsin bleached in any given retina). In WT mice, the response to low intensity generally showed a positive peak at the onset of the light pulse (ON P1), followed by an undershoot that outlasted the step and then returned to baseline (Figure 4B). Responses to higher intensities showed two successive ON response peaks (ON P1 and ON P2), which developed into an undershoot during the pulse. The pulse offset triggered a positive peak (OFF P) that was often followed by an undershoot before returning to baseline. The peaks are consistent with elevated action potential firing of ON, OFF, and/or ON/OFF RGCs. The biphasic ON peak may reflect ON RGC types whose action potentials exhibit different latencies due to variations in the speed of retinal processing and axonal propagation; it also may reflect response dynamics of individual RGCs. The undershoot may reflect a reduction in firing below the spontaneous rate in darkness, as would be expected from adaptational mechanisms in the retina.

To compare responses between WT and *Opa1^R290Q/+^* mice, we focused on relative rather than absolute response amplitudes to control for variations caused mainly by differences in the seal between the electrode and the nerve.^69^ As illustrated in Figure 4C, we observed that these two genotypes differed at higher light intensities in the relative amplitude of the first and second ON peaks (ON P1 and P2). At the highest intensity tested, the ON P2/P1 ratios were 0.65 ± 0.19 and 0.99 ± 0.083 for WT and *Opa1^R290Q/+^* mice, respectively (Figure 4D, Mean ± SD, n = 7 and 3 retinas, *P* < 10^-5^, effect size of 1.96 by Cohen’s *d*). The larger ratio in *Opa1^R290Q/+^* retinas could be due to a number of factors, including RGCs having slower, more dispersed responses (e.g., from poorer conduction in axonal degeneration), RGCs having more sustained responses (e.g., changes in excitability may accompany degeneration), fewer RGCs with faster kinetics, and/or more RGCs with slower kinetics.^72,73^ Therefore, the altered CAPs in *Opa1^R290Q/+^* retinas are consistent with the observed RGC loss and optic nerve degeneration in these mice.

### *Sarm1* knockout (KO) prevents RGC degeneration in *Opa1^R290Q/+^* mice

SARM1, the key executor of Wallerian Degeneration, has been shown to mediate neurodegeneration in response to mitochondrial damage.^38–40,74^ *Sarm1* KO has also been demonstrated to provide protection against retinal degeneration in glaucoma.^75^ We therefore speculated that SARM1 might also drive RGC degeneration in *Opa1^R290Q/+^* mice and sought to determine whether *Sarm1* KO^76^ could protect against RGC death in *Opa1^R290Q/+^* mice. To this end, we built a large mouse cohort consisting of three genotypes: a) *Opa1^+/+^; Sarm1^-/+^* mice as controls to establish baselines; b) *Opa1^R290Q/+^; Sarm1^-/+^* mice, expected to exhibit RGC degeneration similar to *Opa1^R290Q/+^* single mutants; and c) *Opa1^R290Q/+^; Sarm1^-/-^* mice, in which potential rescue effects could be assessed. We followed these mice from 9 to 21 months of age, dissected retinal whole mounts every 3 months, and stained for the RGC marker RBPMS and the cell death marker phospho-H2Ax, as in Figures 2A-2C.

In 21MO *Opa1^R290Q/+^; Sarm1^-/+^* mice, total RGC numbers were decreased and dying RGCs were more abundant than in controls. Both of these changes were rescued to a remarkable extent by *Sarm1* KO, even at 21 months of age (Figure 5A), and the rescuing effect was apparent throughout the longitudinal study (Figures 5B-C). Even at 15 months, when dying cells are most apparent in the *Opa1^R290Q/+^; Sarm1^-/+^* group, the *Opa1^R290Q/+^; Sarm1^-/-^* group remained indistinguishable from the control group (Figure 5C). We also examined the optic nerve by EM in 12MO animals, when axon degeneration is prominent in *Opa1* mutants (Figures 5D and 2E). Whereas the percentage of degenerating axons in *Opa1^R290Q/+^; Sarm1^-/+^* mice was increased compared to the control, the axonal degeneration was largely rescued by *Sarm1* KO (Figure 5E).

**Figure 5.**
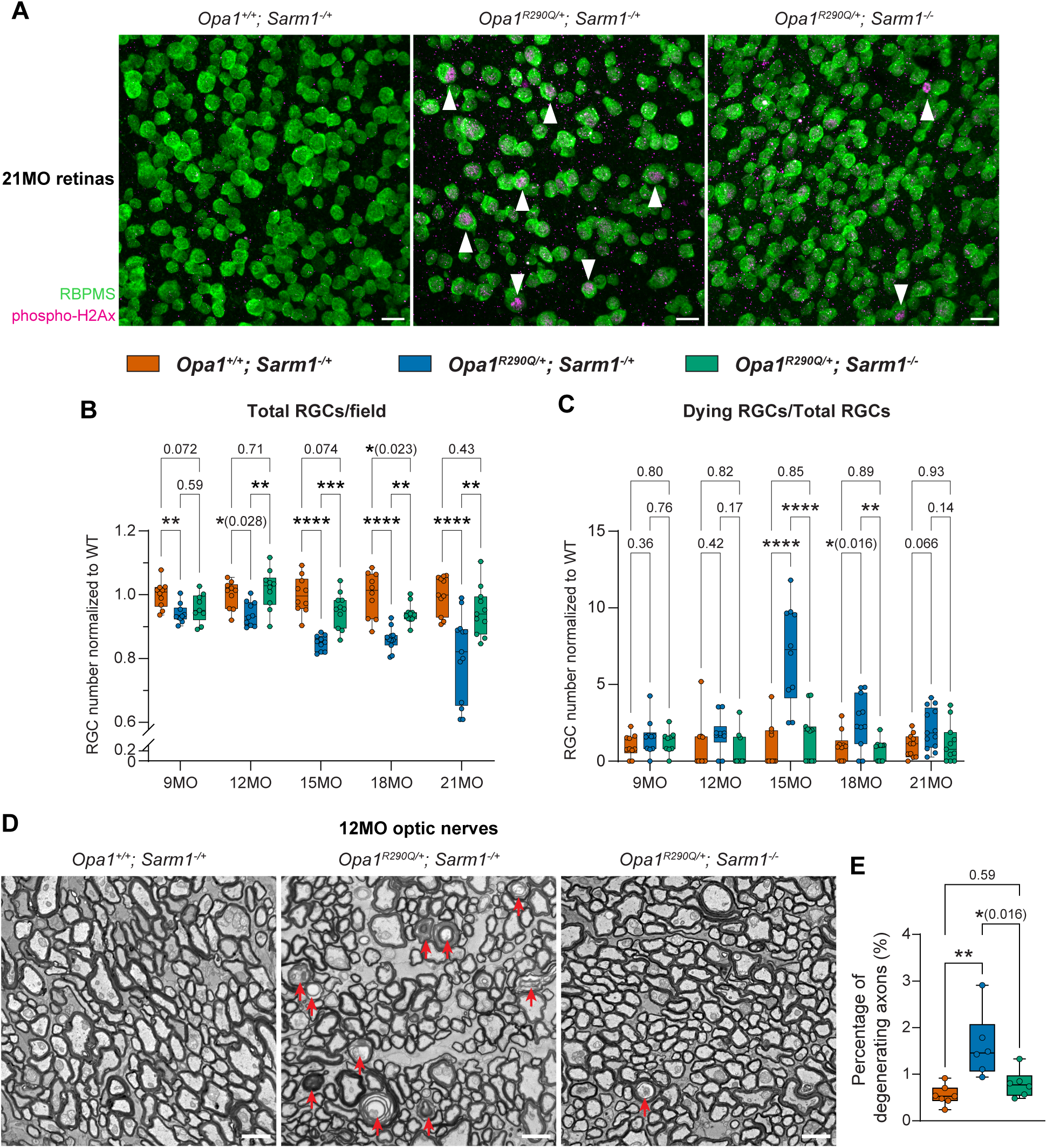
*Sarm1* KO rescues RGC and optic nerve degeneration in *Opa1^R290Q/+^* mice (A) Representative images of retinal whole mounts. Arrowheads indicate phospho-H2Ax- positive dying RGCs. Scale bars = 20 µm. (B) Quantification of total RGCs across ages. Each dot represents one retina. RGC counts were first averaged across the four quadrants of each retina and then normalized to the WT average at each age. N = 10-11 *Opa1^+/+^; Sarm1^-/+^*, 10-13 *Opa1^R290Q/+^; Sarm1^-/+^*, and 9-11 *Opa1^R290Q/+^; Sarm1^-/-^* retinas per age group. (C) Quantification of degenerating RGCs across ages in the same cohort as in (B). Each dot represents one retina. The counts of dying RGCs were averaged across the four quadrants of each retina and divided by the average RGC number of those quadrants. This value was then normalized to the WT average at each age. (D) Representative EM images of cross-sections of optic nerves. Arrows indicate profiles of degenerating RGC axons. Scale bars = 2 µm. (E) Percentage of degenerating RGC axons quantified from EM images in 12MO animals. N = 7 *Opa1^+/+^; Sarm1^-/+^*, 6 *Opa1^R290Q/+^; Sarm1^-/+^*, and 6 *Opa1^R290Q/+^; Sarm1^-/-^* optic nerves. Box plots denote minimum, first quartile, median, third quartile, and maximum values. \**P* < 0.05; \*\**P* < 0.01; \*\*\*\**P* < 0.0001 determined by one-way ANOVA followed by Tukey’s multiple comparisons for each age group.

### *Sarm1* KO rescues the age-dependent decline in RGC function

To examine whether the preservation of RGCs in *Sarm1* KO mice also preserved RGC function, we performed longitudinal ERG and VEP recordings in a separate cohort of mice with the same three genotypes. At all ages examined (4, 12, 15 and 18 months), we observed no differences in the amplitudes of the a-wave and b-wave in flash ERGs (Figures S6A-S6C), confirming that photoreceptors, bipolar cells and Müller glial cells were unaffected by the *Opa1^R290Q/+^* mutation.

At 4 months of age, prior to any signs of RGC degeneration and consistent with normal retinal development in these mice, the three groups did not differ significantly in VEP responses, except for a slightly higher response in the *Opa1^R290Q/+^; Sarm1^-/-^* group compared to the *Opa1^R290Q/+^; Sarm1^-/+^* group (Figures 6A-6B). Starting at 12 months and persisting to 18 months, fVEP and pVEP N1 amplitudes decreased in the *Opa1^R290Q/+^; Sarm1^-/+^* mice. Remarkably, *Sarm1* KO rescued the VEP responses to control levels nearly at all ages examined (Figures 6A-6B). Even at 18 months when the R290Q mutation produced the strongest decrease in this functional assay, the fVEP and pVEP response traces in the *Opa1^R290Q/+^; Sarm1^-/-^* group were indistinguishable from those in the *Opa1^+/+^; Sarm1^-/+^* group (Figures 6C-6D).

**Figure 6.**
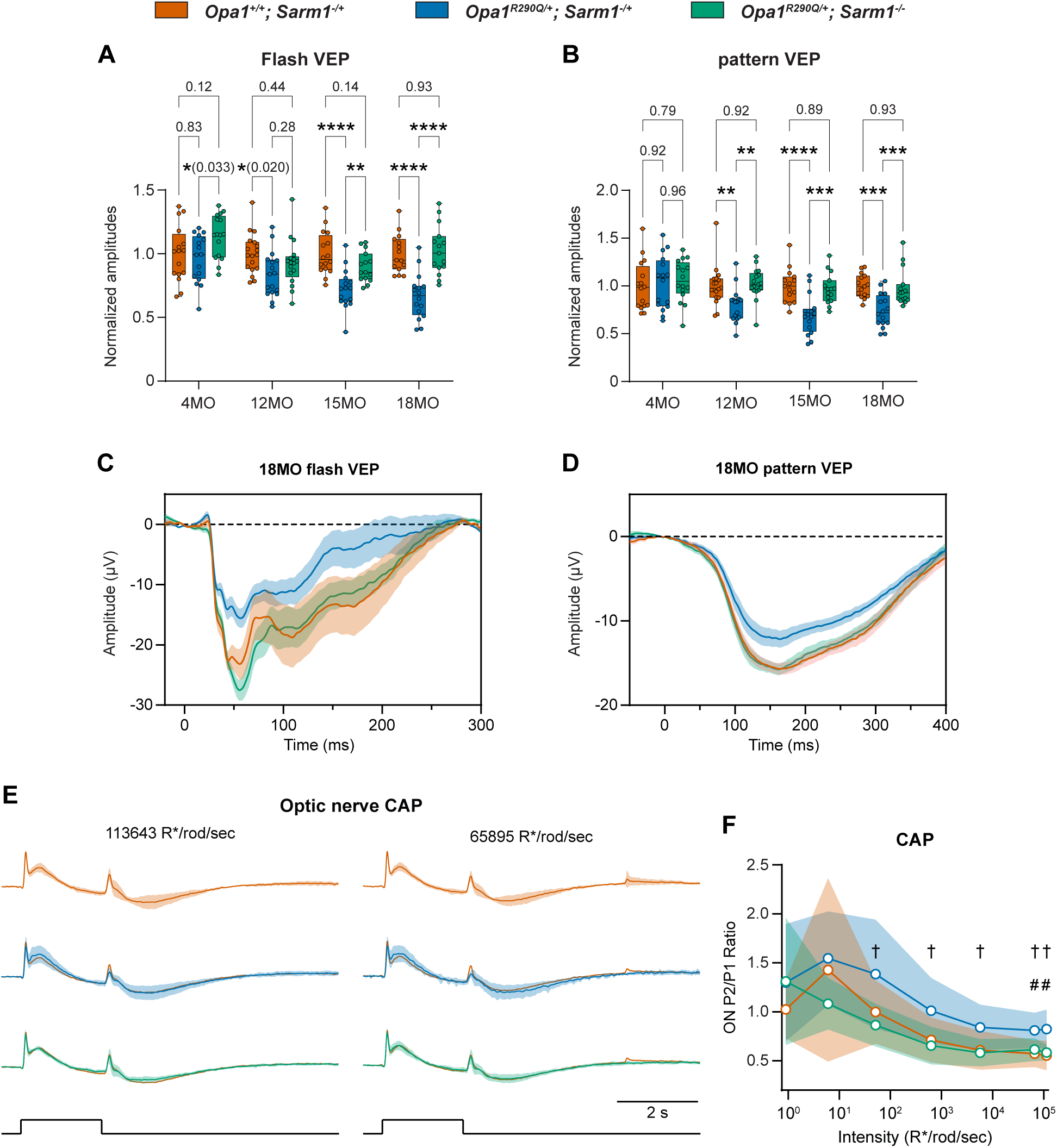
*Sarm1* KO rescues decline in RGC function in *Opa1^R290Q/+^* mice (A-B) Quantification N1 amplitudes of flash VEPs and pattern VEPs measured at the indicated ages of the cohorts. Each dot represents one animal. Amplitudes were normalized to the WT average at each age. N = 15-18 *Opa1^+/+^; Sarm1^-/+^*, 16-17 *Opa1^R290Q/+^; Sarm1^-/+^*, and 15-17 *Opa1^R290Q/+^; Sarm1^-/-^* mice per age group. One-way ANOVA followed by Tukey’s multiple comparisons. Box plots denote minimum, first quartile, median, third quartile, and maximum values. (C-D) Average flash VEP and pattern VEP traces in 18MO animals. Mean ± SEM. (E) Normalized CAP traces from the three genotypes at the two highest light intensities. Mean ± SD. Stimulus monitor at the bottom. The *Opa1^+/+^; Sarm1^-/+^* traces were superimposed onto the other two groups for comparisons. N = 4-5 *Opa1^+/+^; Sarm1^-/+^*, 8-9 *Opa1^R290Q/+^; Sarm1^-/+^*, and 7 *Opa1^R290Q/+^; Sarm1^-/-^* retinas. (F) Ratio of second and first ON peaks (ON P2/P1) as a function of light intensities from (E). Mean ± SD. Mann-Whitney test with bootstrapping (see methods). *P* values for all comparisons are presented in Table S1. \**P* < 0.05; \*\**P* < 0.01; \*\*\**P* < 0.001; \*\*\*\**P* < 0.0001; # marks *P* < 0.05 between *Opa1^+/+^; Sarm1^-/+^* and *Opa1^R290Q/+^; Sarm1^-/+^* retinas; † marks *P* < 0.05 between *Opa1^R290Q/+^; Sarm1^-/+^* and *Opa1^R290Q/+^; Sarm1^-/-^* retinas.

We also recorded CAPs in a randomly chosen subset of this cohort at 19-21 months of age. To enhance signal resolution, we designed and used a new electrode that increases the resistance between the segment of recorded nerve and the bath (see methods). As expected, the ON P2/P1 ratio differed between *Opa1^+/+^; Sarm1^-/+^* and *Opa1^R290Q/+^; Sarm1^-/+^* mice at high intensities (at the highest intensity, the ratio was 0.55 ± 0.15 and 0.82 ± 0.20, respectively; *P* = 0.028, effect size = 1.46, 4 and 8 retinas) (Figure 6E shows the two highest intensities, while Figure S6D shows the lower intensities). The ON P2/P1 ratio was rescued in the *Opa1^R290Q/+^; Sarm1*^-/-^ mice (0.58 ± 0.08, *P* = 0.014, effect size =1.54, 7 and 8 retinas), and the average response in *Opa1^R290Q/+^; Sarm1*^-/-^ mice was indistinguishable from the control (Figures 6E and 6F). Thus, the CAP data support a protective effect from loss of *Sarm1*.

### SARM1 localizes to mitochondrial intermembrane space

Our data suggest a model in which SARM1 becomes activated in the *Opa1* mutant to trigger RGC degeneration. The activation of SARM1 in this ADOA model and by other types of mitochondrial dysfunction, ^38–40^ may be facilitated by the localization of SARM1 to mitochondria. The N-27 amino acids (S27) of SARM1 form a non-canonical mitochondrial targeting sequence capable of lipid binding.^77^ In HEK 293T cells, S27 is sufficient to localize EGFP inside mitochondria.^77,78^ To examine SARM1 localization in neurons, we first overexpressed SARM1- 3×HA in cultured cortical neurons and visualized it with expansion microscopy. The overexpressed SARM1 proteins localized predominantly to mitochondria with no other cellular compartment detectably above background (Figure 7A), consistent with previous reports.^76,77^ We then extracted crude mitochondrial fractions from WT mouse whole brain tissues and examined the localization of endogenous SARM1 with a specific monoclonal anti-SARM1 antibody (Figure S6A).^79^ While SARM1 was abundant in the cytosolic fraction, a significant portion (∼26%) was present in the mitochondrial fraction (Figure 7B). Therefore, at least a fraction of endogenous SARM1 is associated with mitochondria in neurons.

**Figure 7.**
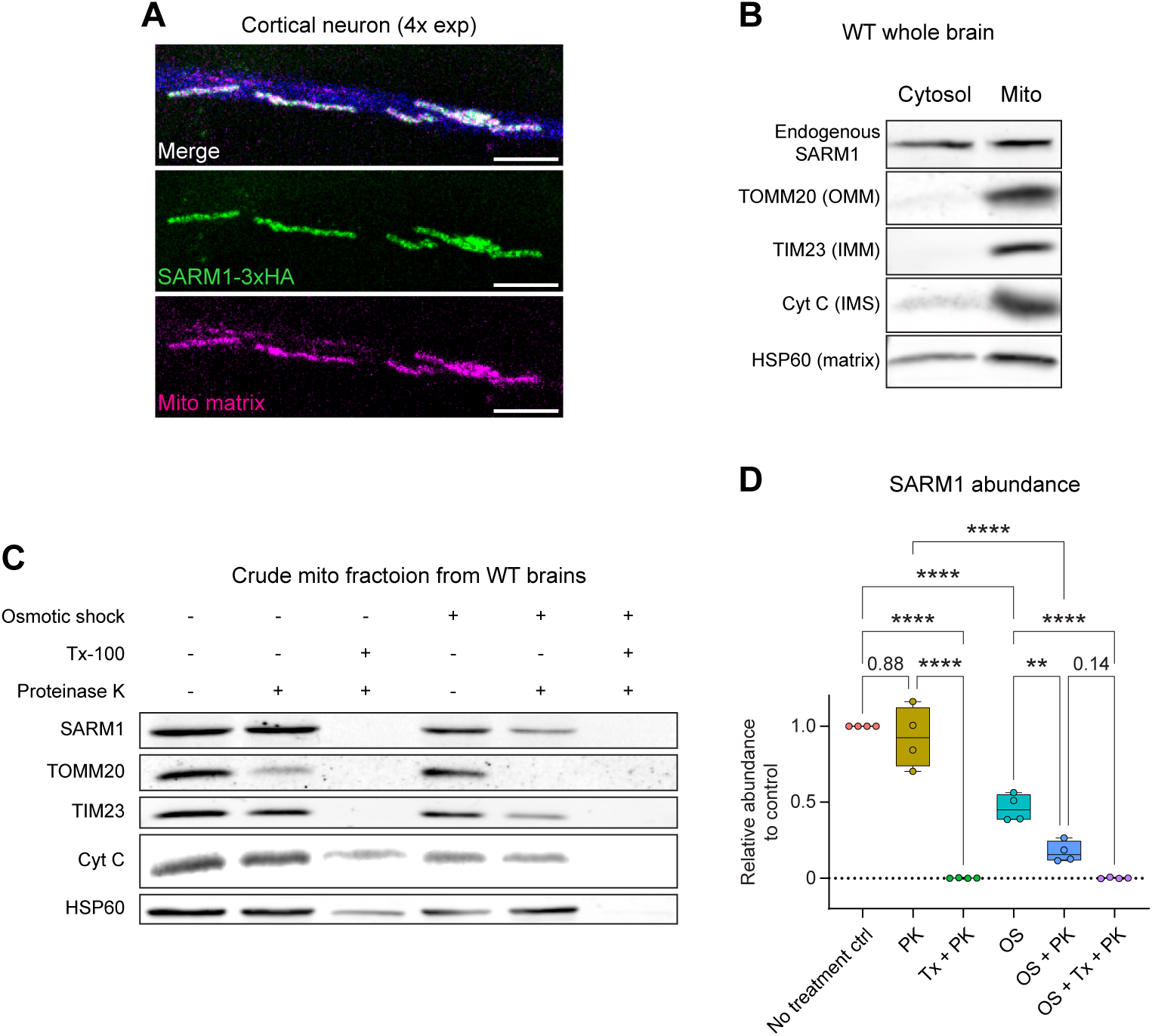
SARM1 localizes to mitochondria IMS (A) Confocal images of overexpressed SARM1 and the mitochondrial matrix marker MitoDsRed in cortical neurons show clear localization of SARM1 to mitochondria. Scale bars = 10 µm. (B) Western blot of SARM1, TOMM20, TIM23, Cytochrome C, and HSP60 in the cytosolic and crude mitochondrial fractions from WT whole brain tissues. OMM, outer mitochondrial membrane; IMM, inner mitochondrial membrane; IMS, intermembrane space. (C) Proteinase K protection assay on crude mitochondria fractions from WT whole brain samples. (D) Quantification of SARM1 abundance across six conditions. N = 4 mice from 4 experiments. Protein levels were normalized to the control condition within each experiment. OS, osmotic shock; PK, Proteinase K. One-way ANOVA followed by Tukey’s multiple comparisons. Box plots denote minimum, first quartile, median, third quartile, and maximum values. \*\**P* < 0.01; \*\*\*\**P* < 0.0001.

Despite SARM1’s mitochondrial targeting sequence, it remains controversial whether endogenous SARM1 in neurons is on the outer membrane (OMM), within the intermembrane space (IMS), associated with the inner membrane (IMM), or inside the matrix.^77,78,80^ To clarify this, we conducted a proteinase K (PK) protection assay on crude mitochondrial fractions extracted from WT whole brains (Figure 7C).^81^ In the untreated control condition, endogenous SARM1, along with TOMM20 (OMM), TIM23 (IMM), Cytochrome C (IMS), and HSP60 (matrix), were all enriched on mitochondria. After a 1-hour PK digestion at room temperature, TOMM20, located on the OMM facing the outside space, was largely degraded. Notably, SARM1 and the other markers were well preserved in the PK-alone condition, indicating that SARM1 is protected by the OMM (Figures 7C-7D and S7B-S7E). Consistent with this, the addition of Triton disrupted mitochondrial membranes, exposing all the marker proteins, including SARM1, to PK treatment.

To further pinpoint SARM1’s localization, we performed osmotic shock (OS) to rupture the OMM and collected the mitoplasts and broken OMM by centrifugation.^82^ As expected, OS treatment significantly reduced Cytochrome C, which was released from the IMS upon OMM rupture (Figures 7C and S7D). Notably, SARM1 levels were reduced by 54% with OS treatment alone (Figure 7D), indicating that much of the mitochondria-localized SARM1 is freely floating in the IMS. OS slightly, but not compellingly, decreased other markers, including TIM23, an integral IMM protein (Figure S7C). When PK was added after OS, both SARM1 and TIM23 were largely degraded, while Cytochrome C and HSP60 levels remained unchanged (Figures 7D and S7C- S7E). The further reduction of SARM1 from OS alone to OS with PK, similar to TIM23, suggests that a fraction of SARM1 is likely anchored to the IMM. These results place SARM1 in two mitochondrial pools: one unanchored pool in the IMS and another pool anchored on the IMM, where OPA1 also resides. Thus, mitochondria-localized SARM1 may occupy a strategic position from which to monitor and respond to electron transport chain damage, such as changes in redox state induced by OPA1 dysfunction.

## Discussion

ADOA is the most common type of inherited optic neuropathy, posing a significant challenge in healthcare. Despite its impact, no therapies are currently available, highlighting a critical unmet need. In this study, we present a novel mouse model of ADOA that, through extensive characterization, reveals robust phenotypes. We identified SARM1, a mitochondria-localized neurodegeneration switch, as a key driver of RGC degeneration in this model. In the *Sarm1* KO background, all the key features of RGC degeneration in the *Opa1^R290Q/+^* mouse were ameliorated, including numbers of surviving RGCs (Figure 5B), counts of dying RGCs (Figure 5C), profiles of degenerating axons (Figure 5E), reductions in visual evoked potentials (Figures 6A-6D), and aberrant compound action potentials in the optic nerve (Figures 6E-6F). The protective effect was long-lasting—even at 21 months of age, the *Opa1^R290Q/+^; Sarm1^-/-^* group remained indistinguishable from the *Opa1^+/+^; Sarm1^-/+^* control group, as measured by both histology and electrophysiology. Our discovery thus opens up exciting possibilities for developing SARM1-targeted therapies for ADOA.

### The *Opa1^R290Q/+^* allele leads to robust mitochondrial and RGC phenotypes

The *Opa1^R290Q/+^* mutation differs from previously generated mouse models in that it is not a simple loss of function allele. The mutation did not reduce overall OPA1 protein levels, though it did alter which protein isoforms were present (Figure S1C). Although some pathogenic *OPA1* mutations, including the most frequent haploinsufficiency allele in human patients, *OPA1^Δ^*^58^*^/+^*, do not cause mitochondrial fragmentation, the *OPA1^R290Q/+^* mutation did.^42^ Therefore, the R290Q mutation appears to be a more severe allele than haploinsufficiency mutations.^41^ In addition, its overexpression induced mitochondrial fragmentation in fibroblasts (Figure S2), where overexpression of the WT protein did not. These findings indicate that the R290Q mutation possesses a dominant negative effect that may be explained by the location of the mutation. Structural studies of the soluble form of OPA1 have shown that it forms a helical lattice on the surface of membranes independent of OPA1 GTPase activity.^44,45^ The R290Q mutation, which falls within the GTPase domain, likely permits the incorporation of the mutant protein into the lattice. If the mutation decreases GTP-stimulated reorganization of the lattice, however, it may ‘poison’ the entire lattice in a manner that a protein-null (haploinsufficient) allele would not.

The dominant negative properties of R290Q may explain why the heterozygous mice exhibit phenotypes across multiple levels, including changes in cristae ultrastructure (Figures 1G-1N) and mitochondrial morphology (Figures 1B-1D), alterations in glutathione redox state and mitochondrial respiration (Figures 1F, S3A, and S3C-D), RGC axon degeneration and cell death (Figure 2), as well as a decline of physiological responses to visual stimuli in the optic nerve (Figure 4) and visual cortex (Figure 3). By following these phenotypes in a longitudinal study, we could describe the progression of the pathology in detail. Previous ADOA mouse models, including those with a Q285stop mutation,^14^ a c.1065+5G splice site mutation,^15^ and a c.2708_2711delTTAG model,^16^ all reflect haploinsufficiency and generally show a milder phenotype. Therefore, the *OPA1^R290Q/+^* mouse model is valuable from a therapeutic development perspective: if a therapy proves effective in addressing the more severe dominant negative scenario, it is likely to be effective in the haploinsufficiency cases as well.

Strong dominant negative *OPA1* mutations have been associated with ADOA plus in some human patients, which includes additional symptoms such as deafness, ataxia, myopathy, peripheral neuropathy and progressive external ophthalmoplegia.^12,13^ However, aside from the RGC phenotypes, the *Opa1^R290Q/+^* mice appear superficially normal and have normal lifespans, though the potential involvement of other organs cannot be excluded.

### Timing and identity of dying RGCs

The relatively short lifespan of the mouse can pose a challenge for modelling a slow neurodegenerative disease. In some cases, mutations that are pathogenic in humans, such as those in the PINK1 and Parkin genes that cause Parkinson’s disease, fail to cause significant neurodegeneration in mice.^83–85^ In patients, ADOA is usually detected within the first decade of life, and progresses slowly over the next two decades or more.^58^ We were fortunate, therefore, to be able to detect strong phenotypes in the *Opa1^R290Q/+^* mouse with RGC death and axon degeneration detectable around 9 months of age, followed by a significant net loss of RGCs and electrophysiological consequences detectable starting at 12 months. The dominant negative properties of the R290Q mutation compared to haploinsufficiency models may contribute to the robustness of the phenotypes. The degeneration progresses slowly such that over 80% of RGCs are still preserved at 18 months, near the end of the normal lifespan of the mice. This observation aligns with the slow progressive nature of ADOA in human patients.^58^ The slow degeneration in the mouse model, however, still provides a substantial window in which to test potential interventions; any such trials should probably be initiated within the first 6 months to effectively ameliorate disease onset.

In the *Opa1^R290Q/+^* mouse, RGC degeneration predominantly occurs between 9 and 15 months, and tapers off by 18 months (Figures 2B and 2E). This pattern might be explained by differential susceptibility of RGC subtypes to mitochondrial damage: certain subtypes may be more vulnerable and degenerate within this window, while others may be more resistant, dying at a slower rate or not at all within the lifespan of the mouse. To date, at least 46 RGC subtypes have been identified in mice, with no single subtype comprising more than 10% of the total RGC population.^86^ These RGC subtypes can be grouped into subclasses with specific molecular markers, such as α-RGCs, T-RGCs, F-RGCs, N-RGCs, T5-RGCs, ipRGCs, and ooDSGCs.^86^ To a large extent, these subclasses appear evolutionarily conserved across species, including humans, though the distribution of each subtype can be different.^87^ It has not been determined in human ADOA patients which RGC subtypes are more vulnerable to degeneration. A future direction beyond the scope of this study will be to screen RGC subtypes and identify those that preferentially undergo degeneration in the *Opa1^R290Q/+^* mouse during aging. Given that the OMR assay did not reveal strong defects in the *Opa1^R290Q/+^* mouse, it is likely that the ON direction-selective RGCs do not undergo significant degeneration.

### SARM1 and Mitochondrial Dysfunction

SARM1 was initially identified as the executioner of Wallerian degeneration and was later shown to play an active role in the pathology of various neurodegenerative conditions, as evidenced by the protection conferred by *Sarm1* knockout (KO) in mouse models.^80,88,89^ Notably, mitochondrial damage activates SARM1, although the precise mechanisms remain elusive.^39^ In this study, we demonstrated that SARM1 is also a key driver of age-related RGC degeneration induced by defective mitochondria in the *Opa1* mutant *in vivo*. SARM1 has a mitochondrial targeting sequence (S27),^77^ but the significance of a mitochondrial pool of SARM1 is unclear; SARM1 can still drive degeneration even without this sequence.^80^ Where SARM1 resides in mitochondria has also been ambiguous. Overexpressed S27-EGFP or SARM1-EGFP localizes inside mitochondria in HEK 293T cells, as confirmed by immunogold labeling and transmission electron microscopy.^77,78^ In contrast, another study reported that overexpressed SARM1-Venus associates peripherally with the mitochondrial outer membrane in cultured rat neurons.^80^ Using immunocytochemistry (ICC), we observed that overexpressed SARM1 predominantly localizes to mitochondria, with no obvious signals in other cellular compartments. Fractionation experiments revealed that a fraction of endogenous SARM1 co-purifies with the mitochondrial fraction, although most remains cytosolic. This discrepancy may arise from the cytosolic portion being too diffuse to detect by ICC or SARM1 leaking from mitochondria during fractionation, leaving the exact size of the mitochondrial pool of SARM1 uncertain. Nevertheless, our proteinase K protection assay clearly indicated that mitochondrial SARM1 localizes to the intermembrane space (IMS) and may also associate with the inner mitochondrial membrane (IMM) (Figure 7C). This localization is intriguing because OPA1 shares a similar localization, raising the possibility of a more direct interaction between the two proteins than previously thought. We speculate that localization of SARM1 in the IMS positions it at an advantageous position to detect defects in the electron transport chain, such as the production of reactive oxygen species.

Is there a specific aspect of mitochondrial damage that activates SARM1? Our data suggests that oxidative stress might play a key role. We observed a significant change in the glutathione redox in both fibroblasts and whole brain metabolites, which is a hallmark of oxidative stress, while ATP production and mtDNA copy number were only minimally affected. Notably, oxidative stress caused by mitochondrial toxins, rather than defective ATP production, has been proposed to robustly activate SARM1 in cell culture models.^39^ This activation likely depends on the loss of NMNAT2, a NAD^+^ synthase that inhibits SARM1.^40^ It would be informative to determine whether SARM1 is activated through a similar mechanism downstream of oxidative stress in ADOA, e.g. by chronic administration of anti-oxidants and examining NMNAT2 levels during aging.

While we demonstrated the comprehensive and enduring protection of *Sarm1* KO in ADOA, it is important to note that the degeneration control group also carried a heterozygous *Sarm1* mutation. Recent studies have shown that loss of one copy of *Sarm1* reduces its activity by half and provides partial protection against neurodegeneration.^90,91^ Since we did not include a littermate control of *Opa1^R290Q/+^; Sarm1^+/+^* in the same cohort, we could not definitively determine whether the heterozygous *Sarm1* mutation also provided protective effects against RGC degeneration in ADOA. However, RGC death in the *Opa1^R290Q/+^; Sarm1^-/+^* mice appears to be delayed compared to the *Opa1^R290Q/+^* single mutants in a separate cohort (Figures 2B and 5C), which supports this possibility. This consideration is particularly relevant for therapy development: small molecule inhibitors or antisense oligonucleotides, which hold great potential as SARM1-targeting therapies, are unlikely to completely inhibit SARM1 activity. Therefore, determining the dose-dependence of RGC degeneration on SARM1 activity will be crucial in the drug development process.

Since the discovery of SARM1,^80,88^ there has been tremendous interest in investigating its involvement in various neurodegenerative diseases. Recent studies have provided compelling evidence that mitochondrial damage is a robust trigger for SARM1 activation and subsequent degeneration.^38–40^ Early research utilized cell culture systems using mitochondrial toxins,^39,40^ while more recent efforts have expanded to *in vivo* models of mitochondrial neurodegeneration, such as Charcot-Marie-Tooth disease type 2A (CMT2A).^38^ Our discovery that *Sarm1* KO suppresses RGC degeneration in the ADOA mouse model further supports the mitochondrial- SARM1 axis as a critical mechanism in mitochondrial neurodegeneration. Although our work does not exclude the involvement of other degenerative pathways, particularly in the context of aging in human ADOA patients, it underscores the critical role of SARM1 in this process. Given this, we propose that SARM1’s role should also be investigated in other types of mitochondrial neurodegenerative disorders, such as Leber Hereditary Optic Neuropathy (LHON), which is caused by mitochondrial DNA mutations and leads to RGC degeneration similar to ADOA.^58,92,93^

## Acknowledgments

We thank Dr. Larry Benowitz, Dr. Chinfei Chen and Dr. Guoli Zhao for constructive discussion. We thank Lala Mkhitaryan for helping with molecular cloning and ordering. We thank Nathaniel Hodgson for training on the OMR assay and the ERG and VEP recording. We thank the Yi-Ping Hsueh lab for providing the mouse monoclonal SARM1 antibody. We thank the Marcia Haigis lab for sharing the Seahorse machine. Figure schematics were generated using BioRender.

We thank several IDDRC Cores supported by NIH P50 HD105351: the Gene Manipulation Core for generating the *Opa1^R290Q/+^* mouse, the Cellular Imaging Core for microscopy services, and the Animal Behavior and Physiology Core for providing the Celeris ERG/VEP platform. We thank the Harvard Medical School Electron Microscopy core for preparing EM samples. We thank the DFCI Metabolomics Core for extracting metabolites from whole brain samples and performing LC-MS. We thank the BIDMC Mass Spectrometry Core for performing metabolomics profiling on fibroblasts metabolites. We are grateful to Sarah Sterling and Jennifer Podgorski at the MIT.nano cryo-EM facility and Zhong Li, Richard Walsh, Megan Mayer, Remya Nair, and Conny Leistner at the Harvard Medical School Cryo-EM Center for Structural Biology.

This work was supported by the Molloy Family Research and Innovation Fund for OPA1 research efforts within the Schwarz lab and the Do lab, NIH R35GM142553 (L.H.C.), the Jane Coffin Childs Fund for Medical Research (M.Y.F.), F.M. Kirby Neurobiology Center Innovation Grants (L.B., C.C., M.T.H.D., and T.L.S.), NIH EY032731 and EY036071 (M.T.H.D.), Helen Hay Whitney Fellowship and HL007901 (S.R.W.), Warren Alpert Distinguished Scholar Award, Lefler Center of Harvard Medical School, and Tommy Fuss Center of Boston Children’s Hospital (P.M.), and the Rosamund Stone Zander Translational Neuroscience Center (RSZ-TNC) postdoctoral fellowship (C.D.).

## Author contributions

C.D., M.T.H.D., and T.L.S. conceptualized the project. C.D. wrote the manuscript and T.L.S. revised the manuscript. C.D. conducted most of the experiments, except where otherwise noted. P.S.N. built and maintained the mouse colony, performed the OMR assay, and conducted part of the EM analyses. J.G. contributed to neuronal culture, expansion microscopy and EM analyses. W.G. contributed to producing the *Opa1^R290Q/+^* mouse. M.Y.F. conducted the Cryo-ET experiments, performed data analyses, and drafted the Cryo-ET results. M.Y.F. and L.H.C. conceived of the Cryo-ET experiment and developed the methodology. S.R.C. conducted the CAP experiments. S.R.W. analyzed the CAP experiments. P.M. conceptualized and developed the CAP technique. M.T.H.D. conceptualized the new CAP electrode, designed the CAP experiments, and drafted the results of these experiments.

## Declaration of interests

No competing interests

## Supplemental information

Document S1. Figures S1-S7 and Table S1 Supplemental figure title and legends

## STAR Methods

### RESOURCE AVAILABILITY

#### Lead contact

Further information and requests for resources and reagents should be directed to and will be fulfilled by the lead contact, Thomas Schwarz.

#### Materials availability

All unique reagents generated in this study are available from the lead contact with a completed materials transfer agreement.

#### Data and code availability

- Cryo-ET data will be deposited at EM databases and will be publicly available as of the date of publication.
- Original western blot images, microscopy data, and CAP data reported in this paper will be shared by the lead contact upon request.
- This paper does not report original code
- Any additional information required to reanalyze the data reported in this paper is available from the lead contact upon request

### EXPERIMENTAL MODEL AND STUDY PARTICIPANT DETAILS

All mouse procedures were approved by the Boston Children’s Hospital (BCH) institutional animal care and use committee (IACUC) and were in accordance with NIH guidelines. Animals were group-housed at the Animal Resources at Children’s Hospital (ARCH) facility and maintained in the environmental conditions recommended by AAALAC. Other procedures involving cell cultures, bacteria, viruses and recombinant DNAs were approved by the Boston Children’s Hospital Institutional Biosafety Committee (BCH IBC).

#### Mouse lines

The *Opa1^R290Q/+^* mice were generated by CRIPSR/Cas9 genome editing on the C57BL/6J background obtained from the Jackson Laboratory (JAX #000664) by the Mouse Gene Manipulation Core at BCH (see method details). All mice were used at the ages specified in the results and figures. All experiments included a balanced mixture of male and female mice in approximately equal numbers.

#### Primary cell cultures

HEK 293T cells (ATCC, CRL-11268) were cultured in Dulbecco’s Modified Eagle Medium (DMEM) with 10% (v/v) fetal bovine serum (FBS) and 1% (v/v) penicillin-streptomycin (P/S) at 37°C with 5% CO2. Primary fibroblasts were isolated as described (see method details) and cultured in DMEM containing 20% FBS and 1% P/S at 37°C with 5% CO2. Both HEK 293T cells and fibroblasts were passaged at 90% confluency. Primary neurons were dissected (see method details) and cultured in Neurobasal (NB) medium supplemented with B27 and Penicillin-Streptomycin-Glutamine (PSG) at 37°C with 5% CO2. The neuronal culture medium was refreshed every two days with a half-volume change.

### METHOD DETAILS

#### Generation of the *Opa1^R290Q/+^* mouse

All animal research and care procedures were approved by the Boston Children’s Hospital IACUC. Mice were maintained on a 12-hour light/dark cycle with food and water provided *ad libitum*. To prepare the ribonucleoprotein complex (RNP), 0.61 pmol each of *Opa1*-R290Q- crRNA and tracrRNA was conjugated and incubated with Cas9 protein (30ng/ul), according to a published protocol^94^. The *Opa1*-R290Q single strand oligonucleotide donor (IDT) was then mixed (10ng/ul) with RNP to prepare the microinjection cocktail. For knock-in mouse generation, the microinjection cocktail was injected into 0.5 dpc embryos harvested after mating C57BL/6J mice (Jackson Laboratory). Post-injection embryos were reimplanted into pseudo-pregnant CD1 foster females (Envigo) and allowed to reach term. Tail snip biopsies were collected from pups at P7 to prepare genomic DNA for screening to identify founders. Genomic DNA was amplified using NEB Q5 Hotstart DNA polymerase with the *Opa1* Genomic PCR screening primers. The PCR products were digested with EcoNI, a site induced by the R290Q mutation. Positive PCR samples were then sequenced with the *Opa1* Genomic PCR sequencing primers. Positive samples were then further validated by sequencing PCR amplicons with the *Opa1* Validation PCR primers. One female founder was obtained, and subsequent genotyping was performed using PCR with the *Opa1* genotyping primers followed by EcoNI digestion. The WT allele produces a 964 bp band, while the R290Q mutation introduces an EcoNI site, resulting in two bands at 356 bp and 608 bp. The female founder was initially outcrossed to wildtype (WT) C57Bl/6J obtained from the Jackson Laboratory (JAX #000664), followed by successive backcrossing to C57Bl/6J. The resulting *Opa1^R290Q/+^* mice were inbred to generate WT and *Opa1^R290Q/+^* littermates. The sequences of the crRNA, HDR template, screening and genotyping primers are listed in Table S1.

#### Isolation of primary fibroblasts

Primary fibroblasts were isolated from mouse embryos from the same litter between embryonic day 15 (E15) and E18. Briefly, the pregnant females were euthanized by CO2, and the embryos were extracted from the uterus. After removing the heads and discarding the viscera, the remaining tissues were finely chopped with a razor blade and digested with 2.5 mL of trypsin for 5 minutes at 37°C. The tissues were then triturated with a P1000 pipette. An additional 2.5 mL of trypsin was added, and the digestion continued for another 5 minutes. 8 mL DMEM containing 20% FBS and penicillin-streptomycin (P/S) was added to terminate digestion. The tissues were then thoroughly triturated with a pipette. After allowing the suspension to settle for 2 minutes, 10 mL of the cell suspension was transferred to a 75cm^2^ tissue culture flask. Fibroblasts were cultured in DMEM containing 20% FBS and P/S, and were passaged upon reaching confluence. Cells were cryopreserved in culture media containing 10% DMSO and stored in liquid nitrogen. Low-passage cells were used for subsequent experiments.

### Isolation of primary neurons

#### Buffers and plates

10X dissection media (DM): 100 mM MgCl2, 10 mM kynurenic acid, 100 mM HEPES in 1X HBSS, pH 7.2.

Papain Solution (10 mL): 10 mL of 1X DM, 200 µL papain, and 8 grains of L-cysteine were mixed and warmed at 37°C until the solution became clear.

Heavy Inhibitor (HI) Solution (10 mL): 8 mL of 1X DM, 80 mg BSA, and 80 mg trypsin inhibitor were combined.

Light Inhibitor (LI) Solution (20 mL): 18 mL of 1X DM was mixed with 2 mL of HI. Culture media: NB supplemented with B27 and PSG

Plates: 12mm #1.5 coverslips were placed in 24-well plates and coated with 3.5 µg/mL laminin and 20 µg/mL poly-L-lysin dissolved in autoclaved ddH2O for 1 hour at 37°C. The plates were then washed three times with water before plating neurons.

#### Dissection and culture

Cortical neurons were isolated from E18-P0 mice from the same litter. Mice were decapitated with scissors and placed in dishes containing 1X DM (10X DM diluted in HBSS), followed by removal of the skin and skull with forceps. The brains were extracted, and meninges were carefully removed. Cortices were dissected and transferred to a 15 mL Falcon tube containing 1 mL of papain solution followed by incubation at 37°C for 5min. The papain solution was then removed, and the tissue was washed with 0.5 mL of LI, repeated twice to thoroughly remove papain. This was followed by a single wash with 0.5 mL of HI. The tissue was then washed three times with 1 mL of culture medium. After the final wash, 1 mL of the culture medium was added, and the tissue was gently triturated ten times with a P1000 pipette. The suspension was allowed to settle for 2 minutes before the cell suspension was transferred to a new tube. Cell density was determined using a hemocytometer, and 100,000 cells in 500 µL were plated per well of a 24-well plate. The media were completely replaced two hours after plating.

### Plasmid construction

Plasmids were constructed using Gibson Assembly or restriction enzyme digestion followed by ligation. For lentiviral constructs, the coding sequences of mouse OPA1, MitoDsRed, mouse Sarm1 were cloned into the pLVX backbone. OPA1 was also cloned into the pEGFP-N1 backbone (Addgene #54767), and the R290Q mutation was introduced using the Q5 site-directed mutagenesis kit. Templates for OPA1, MitoDsRed, Sarm1, and EGFP were obtained from Addgene (plasmid #62845, #174541, #50707, and #54767 respectively). Lentiviral constructs were transformed into NEB stable competent cells, and pEGFP-N1 constructs were transformed into DH5α Competent Cells.

### Lentiviral packaging

HEK 293T cells were seeded into five 15 cm dishes 24 to 48 hours prior to transfection. PEI transfection was done when cells reached 90% confluence. For one dish, 12 µg of psPax2, 5 µg of pMD2.G and 15 µg of transfer plasmid were mixed with 1 mL of DMEM (without phenol red) and 90 µL of PEI. The transfection mixture was then added to the dishes. The following day, the media were replaced with standard HEK culture media. Two days later, the media from all five dishes were combined and filtered through a 0.45 µm filter. The filtered media were centrifuged at 18,000 rpm for 2 hours at 4°C in a Beckman Coulter ultracentrifuge using a SW32Ti rotor.

The supernatant was carefully removed, and the tubes were drained for 5 minutes on a paper towel. The resulting pellet was resuspended in 100 µL of PBS containing 0.001% F68. The resuspended viruses were aliquoted, flash frozen in liquid nitrogen, and stored at -80°C.

### Cell transfection/transduction

#### Fibroblasts transfection

Fibroblasts (60,000 cells per well in 600 µL of culture media) were seeded into 24-well glass bottom plates. The following day, maxipreped plasmids were transfected using the TransIT-LT1 Transfection Reagent. For each well, 0.2–0.5 µg of DNA (OPA1 iso1-WT, OPA1 iso1-R290Q or MitoDsRed) was mixed with 50 µL of Opti-MEM serum-free media and 1.5 µL of the TransIT-LT1 reagent. The mixture was incubated at room temperature for 15 minutes before being added dropwise to each well. Cells were fixed 1 to 2 days post-transfection for immunocytochemistry.

#### Viral transduction

Cultured neurons (100,000 cells in 500 µL of culture media per well) were transduced with lentiviruses encoding MitoDsRed or Sarm1-3×HA. 1.5 to 2 µL of viruses were added per well in 24-well plates on DIV6-7. Neurons were fixed 3-4 days post-transduction for immunocytochemistry and expansion microscopy.

#### Immunocytochemistry (ICC)

For regular ICC, transfected or transduced cells were fixed in 4% PFA for 12 minutes, followed by three PBS washes. Cells were then permeabilized with 0.5% Triton X-100 for 10 minutes, followed by two PBS washes. Blocking was performed using Superblock for 30 minutes. Cells were incubated with primary antibody in Superblock at 4°C overnight. The following day, Cells were washed with PBS three times and incubated with secondary antibody in Superblock for 1 hour at room temperature. After three additional PBS washes, a few drops of Vectashield mounting medium were added, and a #1.5 coverslip was placed over the cells. The cells were imaged on a Zeiss LSM 700 laser scanning confocal microscope with a 63× oil objective.

### Expansion microscopy

#### Staining

Expansion was done following the published glutaraldehyde (GA) method^46^. The staining protocol was similar to the regular ICC protocol with the following modifications: 1) 0.1% GA was included in the fixation medium during the initial fixation step; 2) after fixation, samples were reduced in 10mM sodium borohydride for 30 minutes, followed by three PBS washes before proceeding to permeabilization; 3) after incubation with secondary antibody and subsequent PBS washes, the cells were treated with 0.25% GA for 10 minutes. Following three additional PBS washes, the cells were prepared for expansion.

#### Buffers for expansion

Monomer solution (10 mL): 2.25 mL of 33% w/v Sodium acrylate, 0.625 mL of 40% w/v Acrylamide, 0.75 mL of 2% w/v N, N’-Methylenebisacrylamide, 4 mL of 29.2% w/v Sodium chloride, 1mL of 10X PBS and 1.375 mL of water.

Gelling solution (200 µL): 188 µL of monomer solution, 4 µL of 10% TEMED, 4 µL of 10% Ammonium Persulfate (APS) (add just before gelation) and 4 µL of water.

Digestion buffer (50 mL): 42.5 mL of TAE buffer, 2.5 mL of 10% Triton X-100, 5 mL of 8M guanidine HCl and 1:100 dilution of Proteinase K (20mg/mL) added before use ***Expansion***

Immediately before expansion, the gelling solution was prepared without APS. PBS was aspirated from the wells, and 200 µL of monomer solution was added to each well for equilibration. After 1 minute, the coverslip with cells was transferred to a glass-bottom dish with cell side facing up. APS was then added to the gelling solution, and 100 µL of gelling solution was applied onto the coverslip. A 25 mm coverslip was placed on top of the gelling, creating a coverslip-gel-coverslip sandwich. After 30 minutes of gelation at room temperature, the top coverslip was carefully removed with forceps. The sample was then incubated with 1.5 mL of digestion buffer with Proteinase K for 45 minutes at 37°C. Following digestion, the gel was transferred to a 15 cm dish and allowed to expand in ddH2O for 30 minutes, with the expansion process repeated four times (2 hours total). After expansion, the gel was trimmed into a small piece, which was placed in a glass-bottom dish with cell side facing down. The gel was then sealed with 4 mL of 1% low-meting point agarose gel. The sample was imaged using an inverted Leica TCS SP8 laser scanning confocal microscope.

### Seahorse

24,000 fibroblasts were seeded in each well of a seahorse 96-well plate and grown overnight. The next day, the culture medium was replaced with the Seahorse XF DMEM Medium pH 7.4, supplemented with 10 mM glucose, 2 mM GlutaMax and 1 mM pyruvate before the assay.

Seahorse was performed on the XFe96/XF96 Analyzer (Agilent) using the default mito stress protocol, following the manufacturer’s instructions. 1.5 µM Oligomycin, 2µM FCCP and 0.5 µM Rotenone and Antimycin A were added sequentially. After the seahorse assay, 80 µL of RIPA buffer with protease inhibitors was added to the wells to lyse the cells. Protein concentration was then measured using the BCA assay kit. Oxygen consumption rates were normalized by total protein levels.

### Cryo-Electron Tomography

#### Sample preparation

Fibroblasts were detached from flasks and counted manually. Quantifoil Au 200 mesh holey carbon R2/2 grids were glow discharged for 90 s at 15 mA using a PELCO easiGlow glow discharge system (Ted 44 Pella). At room temperature, 3 µL of sample at a concentration of 1*10^6^ cells/mL were deposited onto grids. After 30 seconds, grids were back-blotted with filter paper for 15 seconds before being plunged into liquid ethane. Subsequent grid handling and transfers were done under liquid nitrogen conditions. Grids were clipped using cryo-FIB autogrids (Thermo Fisher Scientific).

#### Preparation of lamellas for cryo-ET

Grids were loaded two at a time into an Aquilos2 (TFS). After loading, grids were sputter coated with inorganic platinum in a cryo-FIB chamber, then using a GIS were coated in organometallic platinum and then grids were sputter coated with inorganic platinum again to protect and prevent uneven thinning of the sample. Lamella were milled using two rectangular patterns with a gallium ion-beam. Trenches were milled to create macro-expansion joints to improve the stability of the lamella. Tilt angles used for milling ranged from 7°-10°. Lamellas were milled at decreasing ion beam currents from 1 nA to 10 pA resulting in lamellae that were 150-200 nm thick.

#### Cryo-ET data collection and processing

Data were collected using an automated data collection software, Thermo Fisher Scientific Tomography 4, on a FEI Titan Krios equipped with an energy filter (20 eV slit width) at 300 keV and a Gatan K3 direct detector. Images were collected using a defocus range of -3.5 to 4.5 µm using super resolution mode at a calibrated pixel size of 1.375 Å/pixel. Tilt range and dose rate varied depending on prepared lamella, but, generally, collections ranged from -65° to 65° in 2° step increments following the Hagen dose-symmetric scheme.^95^ Using counting mode, 910 to 960 ms tilt images were collected with a frame rate of 152-160 ms and dosage of 0.556-0.602 e/Å per frame accumulated to a target total dose of 170-200 e/Å. Acquired tilted movies were motion corrected and Fourier cropped to a pixel size of 2.75Å/pixel using Relion v.5.0b- 3_cu11.8^96^ to generate dose-weighted micrographs. These micrographs were subjected to contrast transfer function (CTF) refinement (CTFFIND4^97^) and tilt-series were aligned using AreTomo.^98^ CTF-corrected tomograms were reconstructed in Relion5 and denoised with either IsoNET^99^ or cryoCARE.^100,101^ Summed projection images of cryo-tomogram slices were performed in *Dynamo*.^102^

#### Segmentation of tomograms

For segmentation purposes, tomograms were first denoised with cryoCARE^100,101^ and then membranes were boosted using with IsoNet.^99^ In general, the following inputs were used for IsoNet: snrfalloff 0.7; deconvstrength 0.7; density_percentage 50; std_percentage 50.

Membranes were then segmented with Membrain-seg^103^ using the provided pretrained model. Segmentations were cleaned up and membrane labels were assigned in Amira (TFS).

#### Quantification of cryo-electron tomograms

*Quantitative and qualitative analysis of cristae morphology:* Cristae were classified as lamellar (sheet or flat), tubular (straw), invagination (short crista with a CJ), ring, globular (balloon), and undetermined (shape could not be determined).

*Analysis of membrane architecture:* Segmentations of wild-type (n=19) and mutant (n=22) mitochondria from Membrain-seg^103^ with assigned labels were used as input for surface morphometrics^55^. Features extracted were OMM-IMM distance, cristae angle relative to the OMM, and membrane curvedness.

### Tissue preparation

Mice at the desired age were euthanized by isoflurane overdose in a bell jar. Transcardial perfusion was immediately performed using a peristaltic pump. Mice were perfused with 25 mL of ice-chilled PBS, followed by 25 mL of 4% PFA in PBS for retinal histology or 2.5% formaldehyde and 2.5% glutaraldehyde in 0.1M sodium cacodylate (pH 7.4) for electron microscopy (EM) at a low rate of 5 mL/min. For retinal histology, the eyeballs were enucleated, a hole was made in the center of the cornea with a needle, and the eyeballs were post-fixed with 4% PFA for one hour before retinal dissection. For EM, the skull and brain were removed, and the optic nerves were dissected and postfixed in 2.5% formaldehyde and 2.5% glutaraldehyde in 0.1M sodium cacodylate (pH 7.4) overnight before proceeding to EM processing.

### Retinal histology

Fixed eyeballs were transferred to a dish containing TBS. The cornea was removed with dissection scissors, and the lens and vitreous were discarded. The sclera was then peeled away from the retina. Four evenly spaced cuts were made on the isolated retina to allow it to flatten.

The retinas were blocked overnight in a 96 well plate with TBS (pH 7.6) containing 5% normal donkey serum (NDS), 1% BSA and 1% Triton X100. The next day, the blocking solution was replaced with the primary antibody solution containing 5% NDS, 0.2% BSA, 0.5% Triton X-100, guinea pig anti-RBPMS antibody (Novus Biologicals or PhosphoSolutions), and rabbit anti-phospho-H2Ax (Cell Signaling) in TBS. The retinas were incubated with the primary solution for three days on a shaker at 4°C. After incubation, the retinas were washed four times with TBS for 30 minutes each. The retinas were then incubated overnight with the secondary antibody solution, which had the same buffer composition as the primary antibody solution, and included Hoechst, Alexa 488-conjugated donkey anti-guinea pig antibody (Jackson ImmunoResearch) and Alexa 568-conjugated goat anti-rabbit antibody (Invitrogen). The following day, the retinas were washed four times with TBS, mounted on slides with 50 µL of Vectashield mounting media, covered by a #1.5 cover glass, and sealed with nail polish. The retina samples were imaged using a Zeiss LSM 700 laser scanning confocal microscope with a 25× oil objective. For each retina, z stacks of images (256 µm x 256 µm) were captured from each quadrant approximately 1.2 mm from the optic nerve head.

### Electron microscopy of optic nerves

Mice were perfused as previously described with 2.5% formaldehyde and 2.5% glutaraldehyde in 0.1M sodium cacodylate (pH 7.4). The optic nerves, prior to the chiasm, were dissected, cut into two pieces, and post-fixed overnight in the same fixation buffer. The next day, the optic nerves were processed at the EM core at Harvard Medical School. The optic nerves were washed in 0.1M cacodylate buffer and post-fixed with 1% Osmium tetroxide (OsO4)/1.5% potassium ferrocyanide (KFeCN6) for 1 hour, followed by two washes in water. The samples were then washed once in 50mM maleate buffer (pH 5.15, MB) and incubated in 1% uranyl acetate in MB for 1hr. After incubation, the samples were washed once in MB, followed by two washes in water, and subsequently dehydrated in graded alcohols (10 minutes each; 50%, 70%, 90%, 2 X 100%). The samples were then incubated in propylene oxide for 1 hour and infiltrated overnight in a 1:1 mixture of propylene oxide and TAAB Epon (TAAB Laboratories Equipment Ltd). The following day, the samples were embedded in TAAB Epon and polymerized at 60°C for 48 hours.

Ultrathin sections (∼80nm) were cut using a Reichert Ultracut-S microtome, mounted onto copper grids, stained with lead citrate, and examined with a JEOL 1200EX transmission electron microscope. Nine fields were imaged for each cross-section at 2,000X magnification (33.9 µm x 22.2 µm). 10,000X mag images were also taken to show axon degeneration in detail. Images were recorded with an AMT 2k CCD camera and saved as TIFF files.

### ERG/VEP recording

Recordings were performed using the Celeris system (Diagnosys) in a dark room. Mice at the desired age were dark-adapted overnight. The next day, mice were deeply anesthetized with intraperitoneal (IP) injections of 100-120mg/kg ketamine IP and 10 mg/kg xylazine. The anesthetized mouse was placed on the platform heated to 37.4°C, gently stretched to its full length, with its head positioned on a stand. Care was taken to ensure that the mouse’s head remained still during breathing.

For flash ERG and flash VEP, a flash LED stimulator with a built-in Ag/AgCl electrode was placed in close contact with each eye of the animal, and eye gel was applied as a lubricant. A platinum recording electrode for VEP was inserted through the skin and placed subcutaneously along the midline above the visual cortex. A reference electrode for VEP was inserted into the snout, and a grounding electrode was placed subcutaneously near the tail. ERG and fVEP were recorded simultaneously, alternating between eyes, with one electrode serving as the recording electrode and the other as the reference electrode. The recording parameters were as follows: 1 cd·s/m^2^ luminance, 5-second intervals, 60 sweeps per run, a 20-ms baseline, a 0.125-300 Hz bandpass filter for ERG, and a 1-100 Hz bandpass filter for fVEP. The ERG a-wave marker was placed at the lowest peak between 10-40 ms, and the b-wave marker at highest peak between 25-85 ms. The fVEP N1 marker was placed at the lowest peak between 30-50 ms.

Pattern VEP was recorded only from the right eye of each animal. A pattern stimulator with a built-in electrode was placed on the right eye, while the other setups remained the same as for flash recording. The protocol parameters were as follows: horizontal bars at 0.0589 cycles per degree, 340 mm viewing distance, 100% contrast, a view angle of 61°H and 48°V, 2 reversals per sec, 400 ms recording time with a 50 ms pre-stimulus baseline, 450 sweeps per result, 2 results per run, 50 cd/m^2^ luminance, and a 1-100 Hz bandpass filter for pVEP.

### Optomotor reflex assay

The optomotor reflex (OMR) was assessed using the qOMR system (PhenoSys). The setup consisted of a virtual cylinder displaying vertical sine wave gratings, projected in two-dimensional coordinate space on computer monitors arranged in a quadrangle around the testing arena, which was enclosed in a soundproof box. A video camera was positioned directly above the animal to record its behavior. When a rotating grating perceptible to the mouse was projected onto the cylinder wall, the mouse tracked the grating with reflexive head movements aligned with the rotation. The OmrStudio software automatically scored whether animals tracked the cylinder. The OMR index was calculated as the number of frames with correct tracking divided by the number of frames in which the head moved in the opposite direction to the gratings.

For the visual acuity test, moving sine wave gratings were presented at a fixed contrast of 100% and a fixed speed of 12 deg/sec, with varying spatial frequencies (0.025, 0.05, 0.1, 0.15, 0.2, 0.25, 0.3, 0.35, 0.4, 0.45, 0.5, 0.6, and 0.9 cpd), with the direction of movement changing every 5 seconds. Four sets of stimuli were used, differing only in the order of the spatial frequencies. The animal’s tracking behavior was assessed for 60 seconds at each spatial frequency, with a homogeneous gray stimulus with the same mean luminance for 10 seconds between gratings. These short testing epochs minimized the likelihood of the mouse adapting to the stimulus and confirmed that each animal was capable of tracking when a salient stimulus was present. Each mouse was tested by two of the four stimulus sets on day 1 and the remaining two sets on day 2. The average response to each spatial frequency across the four tests was calculated and plotted. The detection threshold was set at 30% of the maximal OMR index above 1. Visual acuity was determined as the intersection between the response curve with the threshold.

For contrast sensitivity testing, a single stimulus pattern was used: sine wave gratings fixed at 0.2 cpd and 12 deg/sec, with decreasing contrasts (1-0.1, in 0.1 steps). Similar to the visual acuity test, the animal’s tracking behavior was assessed for 60 seconds while the moving gratings were displayed, followed by a homogeneous gray stimulus of the same mean luminance for 10 seconds. Each mouse was tested twice on day 1 and twice again on day 2. The average response to each contrast across the four tests was calculated and plotted. The detection threshold was set at 30% of the maximal OMR index above 1. Contrast sensitivity was then determined as the intersection between the response curve with the threshold.

### Compound action potential recording

#### Animals

Mice were dark-adapted for 1.5 hours to overnight. Both males and females were used, with ages ranging from P606 to P658 for WT and *Opa1^R290Q/+^* mice, and from P580 to P653 for *OPA1^+/+^; Sarm1^-/-^*, *Opa1^R290Q/+^; Sarm1^-/+^*, and *Opa1^R290Q/+^; Sarm1*^-/-^ mice. We analyzed 5-9 retinas from 3-6 mice of each genotype. For initial development and validation of the CAP recording method, including the CAP’s sensitivity to synaptic antagonists and tetrodotoxin, animals of various genotypes were used. No overt variation with circadian time, sex, or age was observed.

#### Electrophysiology

Following dark adaptation, mice were anesthetized with Avertin. Each retina and its attached optic nerve (severed at the optic chiasm) was then carefully dissected from associated tissues in Ames’ medium (United States Biological or Sigma-Aldrich, supplemented with NaHCO3 and equilibrated with 95% O2/5% CO2). The vitreous humor was preserved, with only debris (e.g., retinal pigment epithelium) removed to minimize the chance of mechanically damaging the retina and RGC axons. The optic nerve’s dural sheath was also preserved. The retina was placed on the untreated glass surface of the recording chamber, and was superfused continuously with Ames’ medium (∼5-8 ml/min, 23 °C). The tissue was visualized on an upright microscope using infrared transillumination (850- or 940-nm center wavelength and 30-nm width at half-maximum) and differential interference contrast optics.

The recording electrode was a fire-polished, glass capillary (PG10165-4, World Precision Instruments) containing an AgCl pellet. It was back-filled with Ames’ medium from the recording chamber. A minimal length of the optic nerve (<500 µm) was drawn into the electrode to ensure mechanical stability. The nerve was sometimes repositioned in the electrode to optimize signal-to-noise, making the use of bath solution more practical than a dedicated electrode solution.

In the *OPA1; Sarm1* cohort, a new electrode design was used to enhance signal amplitude. Essentially, this entailed ensheathing the suction electrode (now made with 2-000-210 glass, Drummond Scientific) within a shell that could be filled with air. This shell was cut from a glass capillary (TW100-6, World Precision Instruments), fire-polished to size, and connected with the suction electrode at its base with epoxy. The signal electrode was an AgCl pellet inside a compartment continuous with the suction electrode. The ground and reference electrodes were combined in an AgCl pellet in the bath. This air electrode’s larger dimensions required a longer segment of the optic nerve to be drawn in. This design increased the maximum response amplitude by 1.5- to 2-fold across genotypes.

All pharmacological agents were added to the bath. Tetrodotoxin was applied at 300 nM in Ames’ medium. The synaptic antagonists were 3 mM kynurenate, 100 µM picrotoxin, 100 µM D,L-AP4, and 10 µM strychnine in Ames’ medium.^104,105^

A differential amplifier (A-M Systems Model 3000 with headstage) was used for recording. The signal was filtered between 0.1 Hz and 10 kHz. Sampling always met or exceeded the Nyquist minimum.

#### Optical Stimulation

Light from a 75-W xenon arc lamp was filtered to deplete heat while selecting intensity and wavelength, then focused through a 4× objective to produce a spatially uniform disc, centered on the retina. An electromechanical shutter was used to control stimulus timing. Stimuli were measured at the site of the preparation using a calibrated radiometer and spectrometer. 410 nm light was used (10-nm bandpass), which is near the isosbestic point of the mouse visual pigments, and light intensity is expressed as photoisomerized rhodopsin molecules per rod per second (R*/rod/s), using an effective collecting area of 0.5 µm^2^, a wavelength of peak sensitivity for rhodopsin of 500 nm, and Govardovskii’s spectral template.^106^

### Crude mitochondrial preparation

Mice were euthanized with CO2 for 5 minutes. Whole brains were extracted and homogenized on ice using a Dounce homogenizer with 30 strokes in 5 mL of isolation buffer (250mM Sucrose, 20mM HEPES (pH7.4), 2mM EGTA in H2O). The brain homogenates were then drawn into a syringe through an 18G needle and expelled through a 25G needle, repeated 15 times. The homogenates were centrifuged at 2,000 g for 10 minutes at 4°C, and the nuclei and cell debris were discarded. The supernatant was transferred to a new tube and centrifuged again at 2,000 g for 10 minutes. The resulting supernatant (S1) was collected and centrifuged at 10,000 g for 10 minutes. The supernatant (S2), containing the cytosolic fraction, was harvested, and the pellet, representing the crude mitochondrial fraction, was resuspended in 1 mL of isolation buffer. The resuspended mitochondria were transferred to a 1.5 mL Eppendorf tube and centrifuged at 10,000 g for 10 minutes as a wash step. The resulting mitochondrial pellet was then resuspended in 1 mL of isolation buffer to prepare the crude mitochondrial fraction. 25 µL of the resuspended mitochondria or the cytosolic fraction was mixed with 25 µL of RIPA buffer containing protease inhibitors (Millipore Sigma) and incubated for 10 minutes on ice. The lysates were then diluted 1:5 with RIPA buffer and used for BCA analysis to determine protein concentration. The crude mitochondrial preparation was then diluted to 2 mg/mL and aliquoted into six tubes (200 µL each) for the Proteinase K protection assay. The remaining mitochondria were stored in -80°C. The cytosolic fraction was diluted to 1 mg/mL with RIPA buffer and 4X Laemmli buffer, boiled at 95°C for 5 minutes, aliquoted, and stored at -80°C for western blotting.

### Proteinase K protection assay

#### Buffers and reagents

Hypotonic solution: 5 mM Sucrose, 5 mM HEPES (pH7.4), 1 mM EGTA in H2O Hypertonic solution: 750 mM KCl, 80 mM HEPES (pH7.4), 1 mM EGTA in H2O Proteinase K solution (PK): 30ug/mL proteinase K in isolation buffer Triton X-100 and Proteinase K solution (Tx-PK): 1% (v/v) Triton X-100, 30 µg/mL proteinase K in isolation buffer PMSF solution: 200 mM PMSF dissolved in isopropanol

#### Assay

The six tubes containing aliquoted mitochondria (200 µL each) were centrifuged at 16,000 g for 30 minutes at 4°C. The supernatant was discarded, and the mitochondrial pellets in the first three tubes were incubated with 300 µL of the following: 1) isolation buffer; 2) PK solution; or 3) Tx-PK solution, for one hour at room temperature. After incubation, 3 µL of 200 mM PMSF was added to each tube to terminate digestion, followed by the addition of 100 µL of 4X Laemmli buffer. The samples were then boiled at 95°C for 5 minutes, aliquoted, and stored at -80°C for western blotting. For the remaining three tubes, the mitochondrial pellets were resuspended in 200 µL of hypotonic solution and incubated on ice for 15 minutes to induce osmotic shock.

Subsequently, 200 µL of hypertonic solution was added to re-establish isotonic conditions. The samples were then centrifuged at 3,000g for 15min at 4°C to obtain mitoplasts. The mitoplasts were then treated in the same manner as the first three samples.

### Western blotting

Protein samples were run on SDS-PAGE gels in Tris-Glycine buffer (25 mM Tris, 192 mM glycine, 0.1% SDS). Proteins were transferred to 0.4 µm nitrocellulose membranes in transfer buffer (25 mM Tris, 192 mM glycine, 10% methanol) at 300 mA for 90 minutes in a refrigerator. The membranes were trimmed to the desired size and blocked with 5% (w/v) milk in TBST for 30 minutes. Following blocking, the membranes were incubated overnight at 4°C in TBST containing primary antibodies. The following day, the membranes were washed three times with TBST, 10 minutes each, followed by a one-hour incubation at room temperature in TBST containing fluorescent secondary antibodies (LiCor). After three additional 5-minute washes with TBST, the membranes were imaged using the LiCor Odyssey CLx imager.

### Reverse transcription and qPCR of *Opa1* isoforms

Fibroblasts were seeded in 24-well plates at a density of 60,000 cells per well. The next day, RNA was extracted from each well as separate samples using the RNeasy kit (Qiagen). RNA concentrations were measured using a Nanodrop. For cDNA synthesis, 800 ng of RNAs was used with the iScript Reverse Transcription kit (Bio-Rad). Cortical neurons were dissected from mouse brains and seeded at 10,000 cells per well in 24-well plates. On DIV8, RNA was extracted, and 200 ng of RNA was used for cDNA synthesis.

PCR of *Opa1* isoforms was performed using Q5 Hot Start DNA Polymerase, OPA1-specific primers (see Table S1), and 2 µL of cDNA, with an annealing temperature of 67°C, a one-minute extension time, and 28 cycles. PCR of the ActB control was performed with the same cDNA templates, DNA Polymerase, and ActB specific primers, using a 67°C annealing temperature, a one-minute extension time and 32 cycles. The PCR products were resolved on a 1% agarose gel stained with EtBr.

### mtDNA content measurements

WT and *Opa1^R290Q/+^* MEFs were collected from flasks during passaging, and 1.5-2.5 million cells were used for DNA extraction using the DNeasy kit (Qiagen). DNA concentrations were measured with a Nanodrop and diluted to 40 ng/µL. qPCR was performed using the PowerUp™ SYBR™ Green Master Mix (Thermo Fisher Scientific), gene-specific primers (see Table S1), and 120ng DNA templates. Triplicates were used for each gene and sample. CT values were obtained for each sample, and fold changes were calculated using the ΔΔCT method.

### Metabolomics

#### Fibroblasts

<u>Metabolite extraction.</u> Fibroblasts were seeded in 15 cm dishes and cultured until 90% confluence. Prior to extraction, the cells were gently washed once with 5 mL of regular culture media and incubated with 10 mL of culture media for 2 hours. The media was then completely aspirated, and 4 mL of 80% methanol (-80°C) was immediately added. The dishes were then transferred to -80°C on dry ice and incubated for 15 minutes. After incubation, the cells were scraped with cell scrapers, and the cell lysate/methanol mixture was transferred to 15 mL Falcon tubes on dry ice. The lysates were centrifuged at full speed for minutes at 4°C to pellet cell debris and proteins. The supernatant was collected into 50 mL Falcon tubes on dry ice. 500 µL of 80% methanol (-80°C) was added to the 15 mL tube to resuspend the pellet. The mixture was transferred to1.5 mL Eppendorf tubes and centrifuged at full speed for 5 minutes at 4°C. The supernatant was transferred to the 50 mL Falcon tubes on dry ice containing the supernatants collected before. The extraction was repeated one more time and all three extractions were pooled in the 50 mL tube. The samples were then completely dried using a speedVac for 5 hours.

Polar metabolomics profiling was performed at the Beth Israel Deaconess Mass Spectrometry Core as previously described.^47^

#### Brains

<u>Metabolite extraction.</u> Mice were euthanized by CO2 for 5 minutes, after which whole brains were immediately extracted, weighed, and flash-frozen in liquid nitrogen. Metabolites were extracted from bead-homogenized flash-frozen brains using 7 mL of 70% of 75°C ethanol/water. Extraction was repeated once on the cell debris. The extraction solvent was spiked with 0.1% labelled amino acid mix (Cambridge Isotope Laboratories Inc., MSK-A2-1.2). The extracts were dried in a centrifugal vacuum unit, resuspended water and diluted in 80/20 acetonitrile/water with 1% formic acid to final volume of 400 µL for brain samples, and cleaned using a pass- through plate (Ostro Protein Precipitation & Phospholipid removal plate, Waters, SKU 186005518), dried in a centrifugal vacuum unit, and reconstituted in 80/20 acetonitrile/water.

<u>LC/MS analysis.</u> For metabolite analysis, a mass spectrometer (QExactive HF-X) was equipped with HESI II probe and coupled to a Vanquish binary UPLC system (Thermo Fisher Scientific, San Jose, CA). 5 µL of each metabolite extract was injected onto a BEH Z-HILIC column (100 mm, 1.7 µM particle size, 2.1 mm internal diameter, Waters). Mobile phase A was 15 mM ammonium bicarbonate in 90% water and 10% acetonitrile, and mobile phase B was 15 mM ammonium bicarbonate in 95% acetonitrile and 5% water. The column oven was held at 45°C and autosampler at 4°C. The chromatographic gradient was carried out at a flow rate of 0.5 mL/min as follows: 0.75 min initial hold at 95% B; 0.75-3.00 min linear gradient from 95% to 30% B, 1.00 min isocratic hold at 30% B. B was brought back to 95% over 0.50 minutes, after which the column was re-equilibrated under initial conditions. The analysis was performed in negative mode for each sample and gradient. The mass spectrometer was operated in full-scan mode (m/z = 70–1,050), with the spray voltage set to 3 kV, the capillary temperature to 320 °C, and the HESI probe to 300 °C. The sheath gas flow was 50U, aux gas 10U, and the sweep gas was 1U, resolving power was 120,000. tSIM experiments were set up for GSH and GSSG using authentic standards. Water, formic acid, and acetonitrile were purchased from Fisher and were Optima LC/MS grade. Ammonium bicarbonate powder was purchased from Merck, and ethanol from Decon laboratories.

### QUANTIFICATION AND STATISTICAL ANALYSIS

#### Mitochondrial length measurements

Mitochondrial length was measured in Fiji (ImageJ, NIH). Maximum intensity projections were generated from Z stacks of mitochondrial images and analyzed with the Mitochondrial Network Analysis (MiNA)^107^ plugin. Parameters were carefully adjusted to prevent over-fragmentation or over-fusion artifacts. The “Summed branch lengths mean” outputs were extracted and used for quantification.

#### RGC counts

Maximum projections of retinal stacks were generated using Fiji, and quantified in a blind manner. The total number of RGCs per retina was calculated as the average RGC count across the four quadrants. Dying RGCs were identified by phospho-H2Ax positive nuclei, and the percentage of dying RGCs was determined by dividing the average number of dying RGCs by the average total RGC number for that retina. Both the total number of RGCs and the percentage of dying RGCs were normalized to the WT means for each age group and displayed as box plots showing median, quantiles, minimum, maximum, and individual data points. This normalization facilitates a clearer comparison of the RGC degeneration patterns between WT and mutant mice across different ages, accounting for age-specific variations in WT RGC counts. With this normalization, different age groups were treated as independent datasets, and statistical analyses were performed using Mann-Whitney tests for comparisons within each age group containing two genotypes, and one-way ANOVA with Tukey’s multiple comparisons for each age group with three genotypes.

#### EM analyses

EM images were quantified using Fiji. Numbers of total axons and degenerating axons were counted and averaged across the nine fields of each optic nerve section. The percentage of degenerating axons was calculated by dividing the average number of degenerating axons by the average total number of axons for each optic nerve. These values were then normalized to the WT means for each age group, similar to the RGC counts, and presented as box plots.

Mann-Whitney tests were performed within each age group containing two genotypes, while one-way ANOVA with Tukey’s multiple comparisons was used for each age group with three genotypes.

#### ERG/VEP quantifications

ERG/VEP results were exported from the Celeris system as .csv files. The following values were extracted for each animal: a/b wave amplitudes of flash ERGs from both eyes, N1 amplitudes of flash VEPs from both eyes, and N1 amplitudes of pattern VEPs from the right eye. The a/b wave amplitudes of ERGs and N1 amplitudes of fVEPs were averaged between the two eyes for each animal. These values were then normalized to the WT means for each age group and presented as box plots. Mann-Whitney tests were performed within each age group containing two genotypes, while one-way ANOVA with Tukey’s multiple comparisons was used for each age group with three genotypes. For the response traces in 18 MO animals, raw values were averaged across all animals and plotted as Mean ± SEM.

#### Metabolomics data analysis

Target metabolites and internal standards were analyzed using emzed.^108^ Peak areas were normalized by dividing each peak area by the measured biomass in each sample and subsequently by the average of internal standard peak areas detected in the data file.

Normalized peak areas were used for quantification.

#### CAP analyses

Responses from a retina at a given intensity were baseline-subtracted, averaged, and low-pass filtered (10 Hz) prior to analysis. A permutation test was used to estimate the likelihood that observed differences between samples arose by chance. Data values were resampled using a bootstrapping algorithm, randomly assigned to two groups, and compared using the Mann-Whitney U test. Repeating this operation 100,000 times yielded a distribution of U test statistics, to which the U test statistic of the actual datasets was compared. Effect sizes are given as Cohen’s d. Responses with an average Z-score [(response peak - avg baseline)/ baseline SD] of <10 were not used for statistical analysis.

#### Immunoblot quantifications

Western blots were imaged on the Odyssey CLx imager. Protein intensities were quantified using the Image Studio Light software (LiCor). For the mitochondrial PK assay, signals for each protein were normalized to the untreated control. Results were presented as box plots.

Statistical analyses were conducted using one-way ANOVA followed by Tukey’s multiple comparisons.

#### Statistical analyses

Datapoints represent biological replicates. Box plots denote minimum, first quartile, median, third quartile, and maximum values. Statistical analyses were conducted using Prism (v10; GraphPad Software). Specific statistical analyses used are described in figure legends and methods. P-values are listed in graphs and Table S1.

---

**Supplemental Figure 1 - related to Figure 1.**
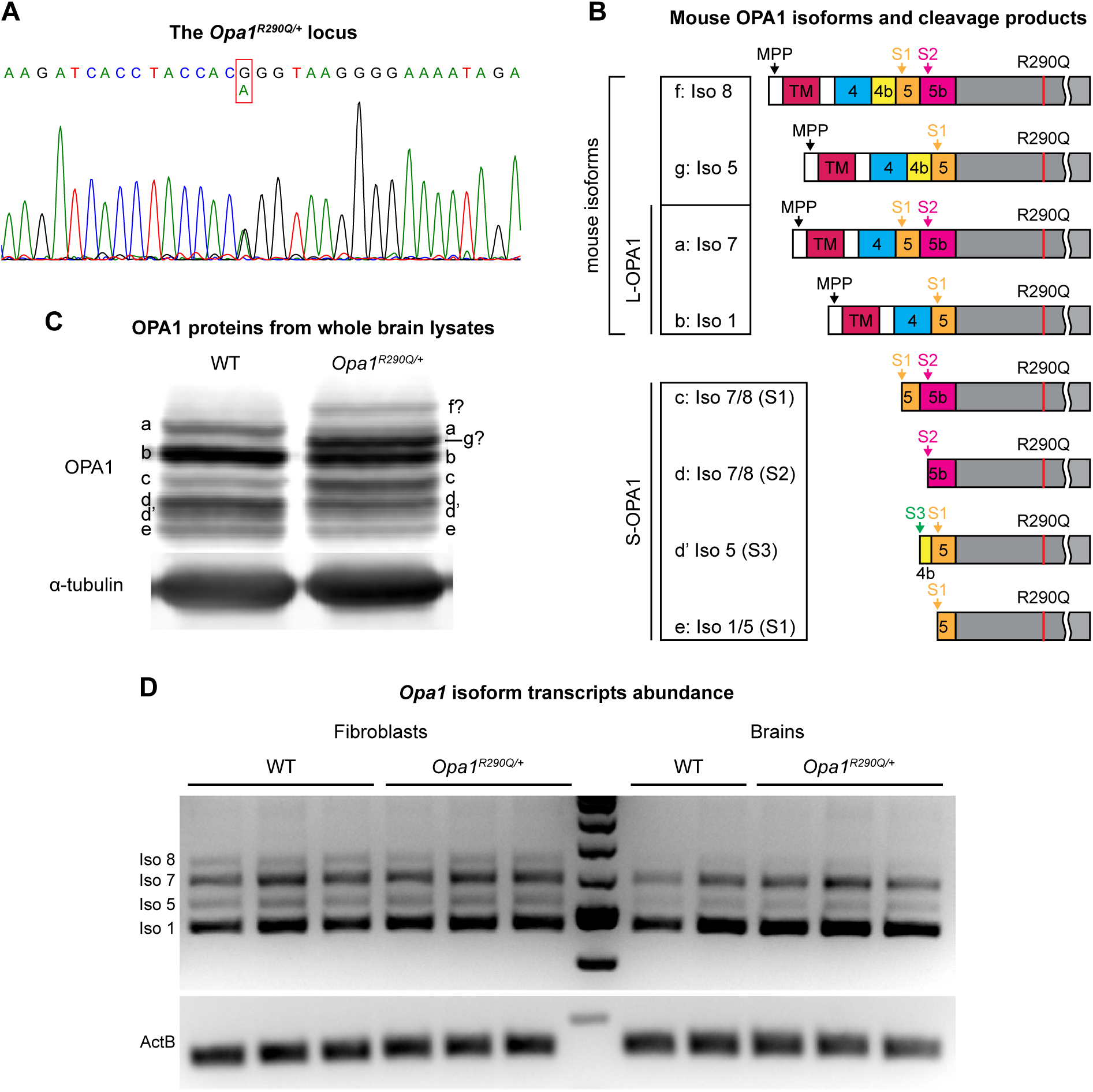
Impact of the R290Q point mutation on *Opa1* isoform abundance and post-translational processing (A) The mouse *Opa1^R290Q/+^* locus and a representative Sanger sequencing result from PCR amplification of genomic DNA. (B) Schematic representation of mouse OPA1 protein isoforms based on previous studies.^19^ The mouse expresses four *Opa1* isoforms. In WT cells, isoforms 5 and 8 are fully cleaved by OMA1 at site S1 and YME1L at site S2, while isoforms 1 and 7 undergo partial cleavage. The R290Q mutation falls within the common region present in all isoforms. MPP, mitochondrial processing peptidase; S1, OMA1 cleavage site; S2, YME1L cleavage site; S3, another YME1L cleavage site. (C) Western blot analysis of OPA1 in brain lysates from post-natal day 0 WT and *Opa1^R290Q/+^* mice. In WT, the OPA1 pattern aligns with previous reports.^19^ In *Opa1^R290Q/+^* mice, two additional longer bands, likely corresponding to full-length isoforms 5 and 8, were observed. The intensity of other bands also changed, though the abundance of total OPA1 proteins remained similar. (D) PCR analysis of OPA1 isoform abundance in genomic DNA extracted from fibroblasts (n = 3 WT and 3 *Opa1^R290Q/+^* passages) and brain tissues (n = 2 WT and 4 *Opa1^R290Q/+^* brains). ActB was used as the control. 4 splicing variants arose via the inclusion or exclusion of exons 4b and 5b and no significant changes due to R290Q were detected.

**Supplemental Figure 2 - related to Figure 1.**
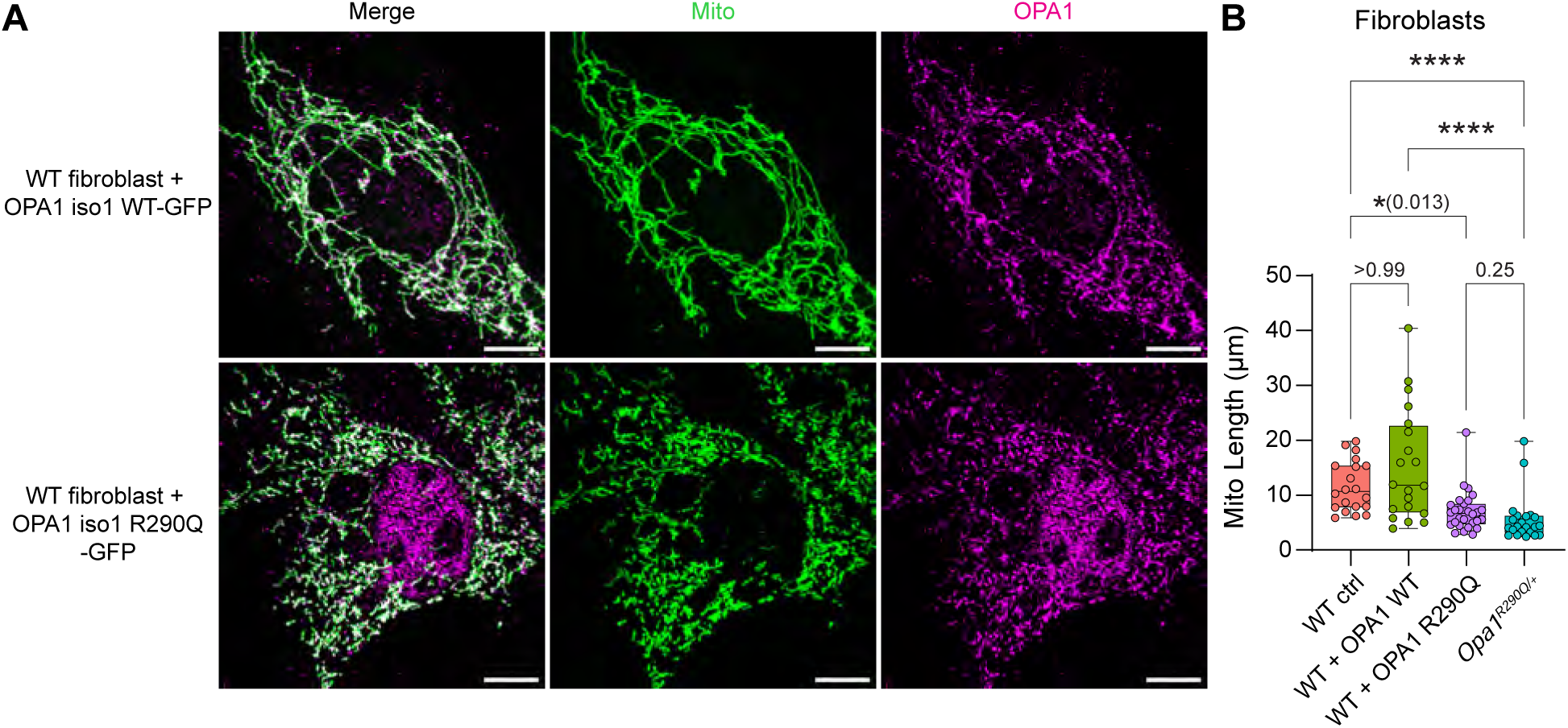
The OPA1 R290Q transgene causes mitochondrial fragmentation upon overexpression (A) Representative images of WT fibroblasts expressing transgenes encoding either OPA1 isoform 1 WT-GFP or OPA1 isoform1 R290Q-GFP. Mitochondria (green) were labeled by expression of MitoDsRed and OPA1 (magenta) was labeled with GFP staining. Scale bars = 10 µm. (B) Quantification of mitochondrial length in fibroblasts overexpressing OPA1 as in (A). Overexpression of OPA1 R290Q, but not WT OPA1, in WT fibroblasts resulted in mitochondrial fragmentation comparable to that observed in heterozygous fibroblasts. N = 20, 20, 28, and 22 cells for each condition, respectively, from one experiment. Kruskal-Wallis test with Dunn’s multiple comparisons. Box plots denote minimum, first quartile, median, third quartile, and maximum values. \**P* < 0.05; \*\*\*\**P* < 0.0001.

**Supplemental Figure 3 - related to Figure 1.**
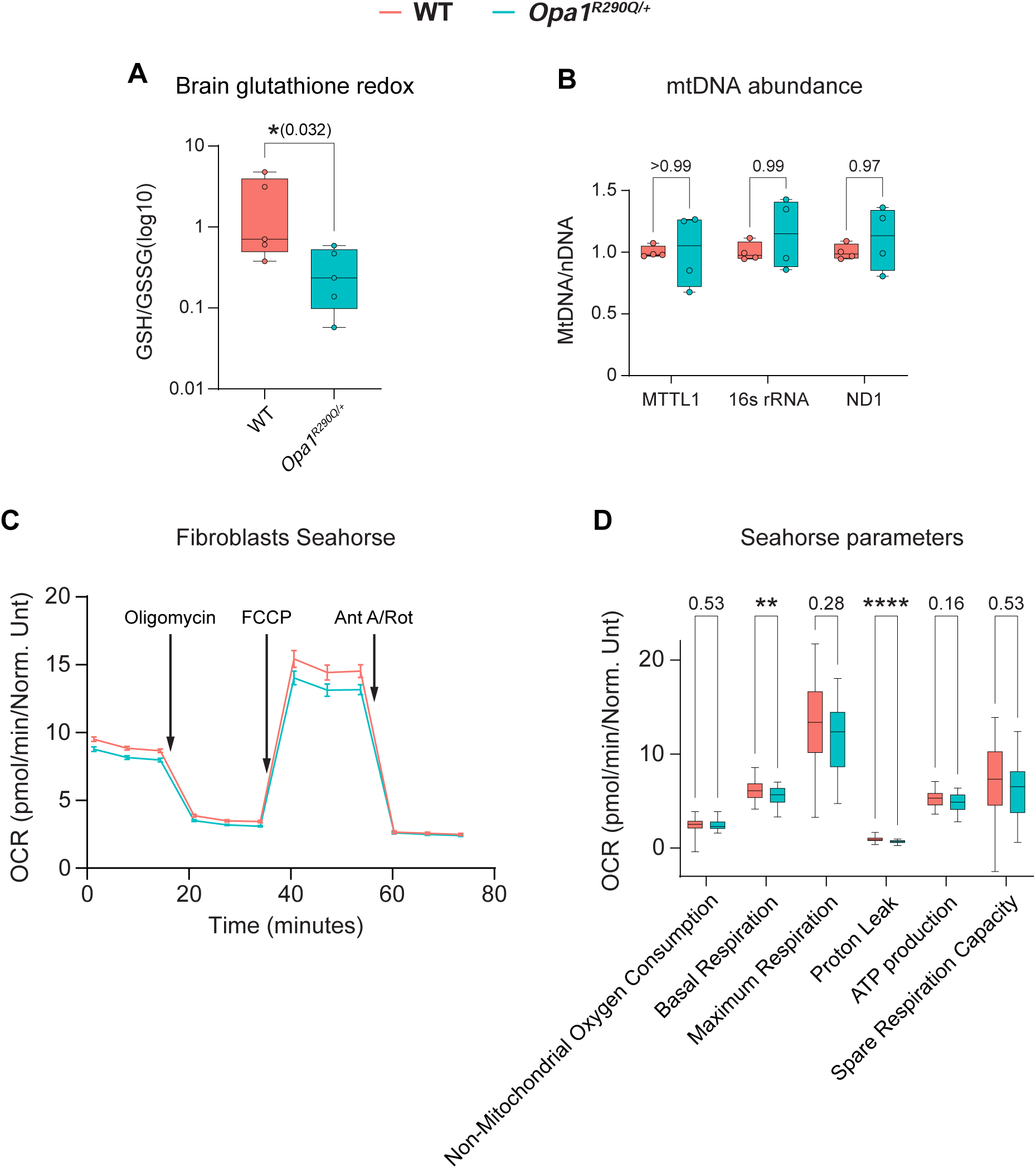
The *Opa1^R290Q/+^* mutation causes oxidative stress but has minimal impact on oxygen consumption and mitochondrial DNA stability in fibroblasts (A) GSH/GSSG ratios in whole brain tissues as measured by LC-MC, as also shown in fibroblasts (Figure 1F). N = 5 WT brains and 5 *Opa1^R290Q/+^* brains from one experiment. Mann-Whitney test. (B) Mitochondrial DNA abundance was measured by qPCR in DNA extracted from fibroblasts. GAPDH DNA was used as the nuclear DNA reference for normalization. N = 4 WT and 4 *Opa1^R290Q/+^* samples from different passages. Multiple Mann-Whitney tests with Holm-Sidak multiple comparisons. (C) Oxygen consumption rates of fibroblasts, as measured by the Seahorse assay. N = 62 WT wells and 58 *Opa1^R290Q/+^* wells from 4 experiments. (D) Parameters of mitochondrial function derived from the measurements in (C). Multiple unpaired t tests with Holm-Sidak multiple comparisons. Box plots denote minimum, first quartile, median, third quartile, and maximum values. \**P* < 0.05; \*\**P* < 0.01; \*\*\*\**P* < 0.0001.

**Supplemental Figure 4 - related to Figure 3.**
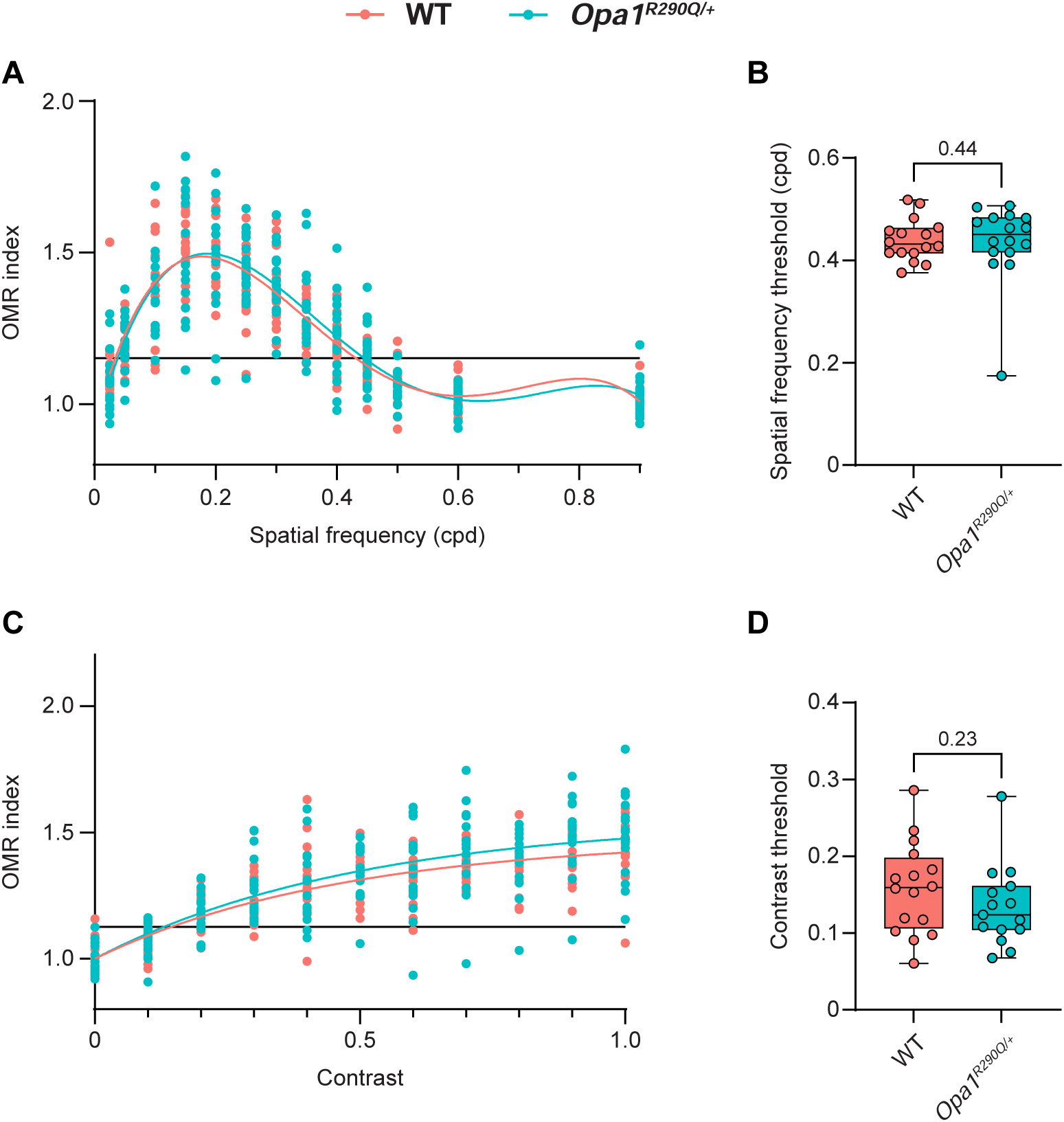
*Opa1^R290Q/+^* mice do not show significant decrease in the optomotor reflex (OMR) (A) OMR index plotted as a function of spatial frequency (cycles per degree). The data was fitted with a 4^th^ degree polynomial function. The horizontal line represents the OMR response threshold, set at 30% of the maximal response above a baseline of 1. (B) Spatial frequency thresholds determined by the second intersection of the horizontal line in (A) with the fitted curve for each animal. N = 16 WT and 16 *Opa1^R290Q/+^* mice. Mann-Whitney test. (C) OMR index plotted as a function of contrast (0 to 1). The data was fitted with a one phase association function. The horizontal line represents the OMR response threshold, set at 30% of the maximal response above a baseline of 1. (D) Contrast thresholds determined by the intersection of the horizontal line in (C) with the fitted curve for each animal. N = 16 WT and 15 *Opa1^R290Q/+^* mice. Mann-Whitney test.

**Supplemental Figure 5 - related to Figure 4.**
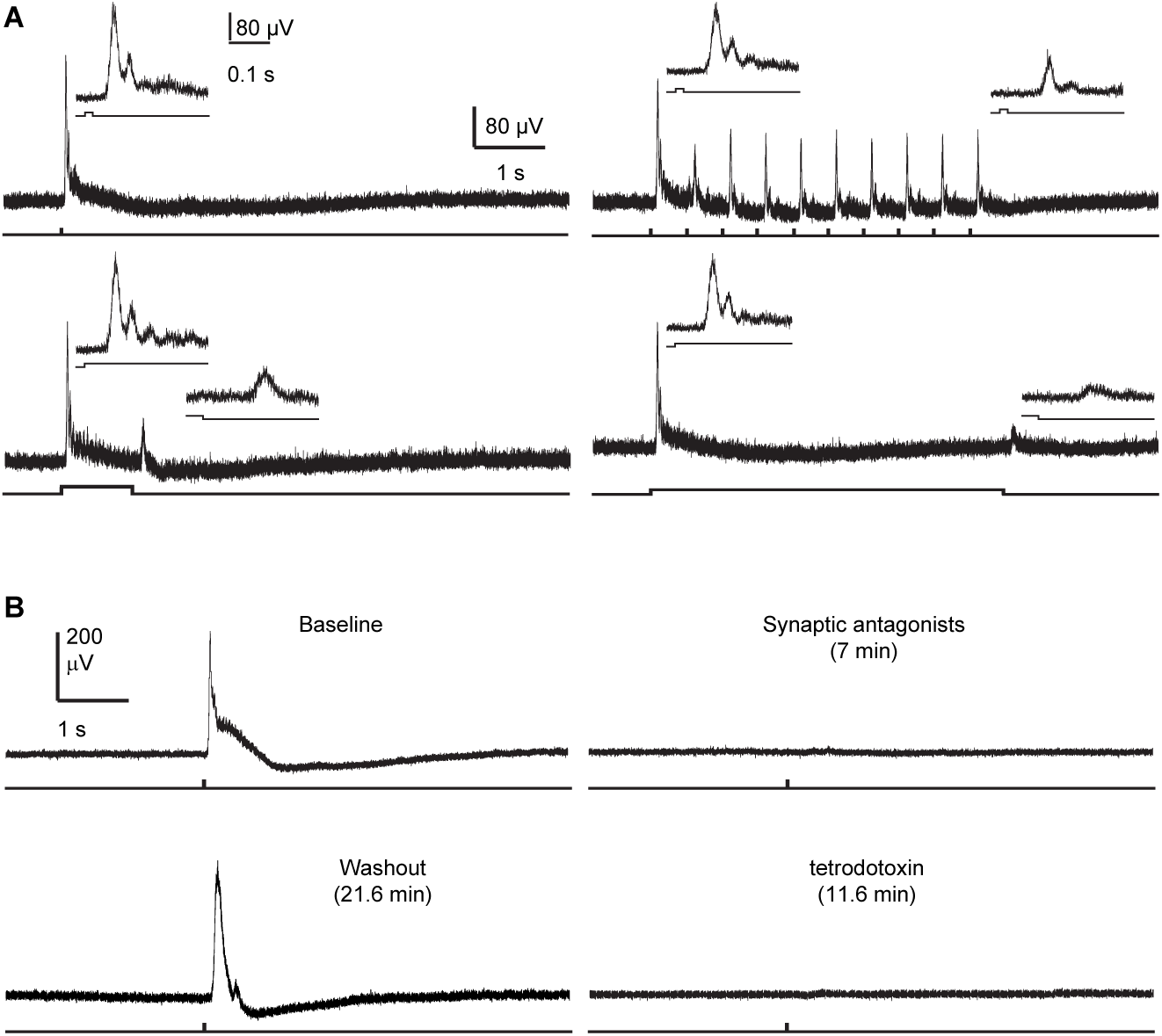
Basic features of the compound action potential (A) Responses to flashes and steps of light. (B) Block by pharmacological antagonists of synaptic transmission (top, 3 mM kynurenate, 100 µM picrotoxin, 100 µM D,L-AP4, and 10 µM strychnine) and action potentials (bottom). These effects were reversible (not shown). Stimulus monitors are shown at the bottom.

**Supplemental Figure 6 - related to Figure 6.**
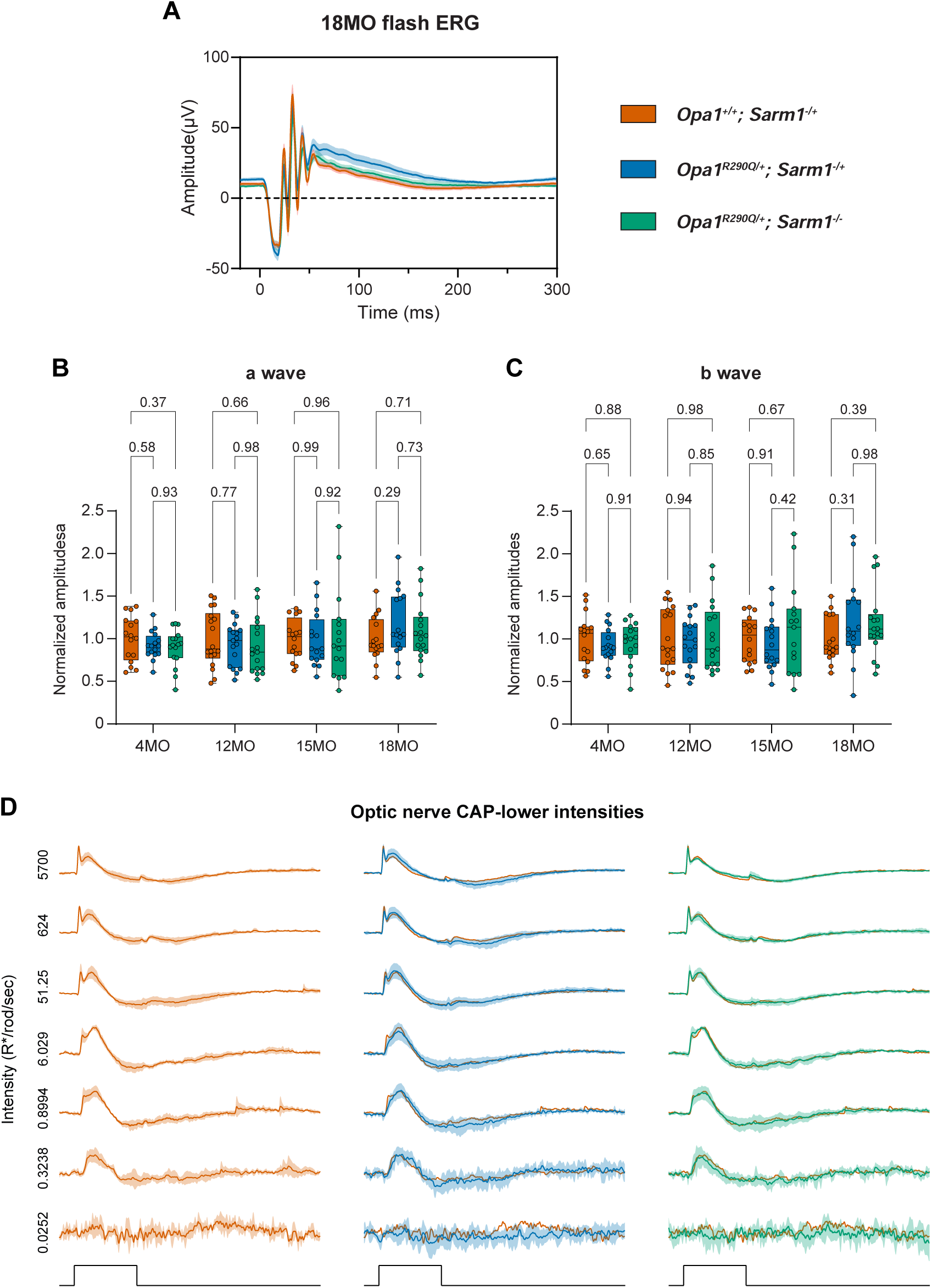
Flash ERG responses are not appreciably impacted by either the *OPA1^R290Q/+^* mutation or *Sarm1* KO (A) Average flash ERG traces in 18MO animals of the indicated genotypes. N = 16 *Opa1^+/+^; Sarm1^-/+^*, 16 *Opa1^R290Q/+^; Sarm1^-/+^*, and 17 *Opa1^R290Q/+^; Sarm1^-/-^* mice. Mean ± SEM. (B-C) Amplitudes of a-wave (B) and b-wave (C) of flash ERG at the indicated ages of the cohorts. Each dot represents one animal. Amplitudes were normalized to WT average at each age. N = 16-18 *Opa1^+/+^; Sarm1^-/+^*, 16-17 *Opa1^R290Q/+^; Sarm1^-/+^*, and 15-17 *Opa1^R290Q/+^; Sarm1^-/-^* mice per age group. One-way ANOVA followed by Tukey’s multiple comparisons for each age group. Box plots denote minimum, first quartile, median, third quartile, and maximum values. (D) Similar to Figure 6E, but at lower light intensities. Mean ± SD. Stimulus monitor at the bottom. The *Opa1^+/+^; Sarm1^-/+^* traces were superimposed onto the other two groups for comparisons. For the lowest two intensities, distinct peaks were poorly discriminated (average Z-score < 10) and were not included in analyses.

**Supplemental Figure 7- related to Figure 7.**
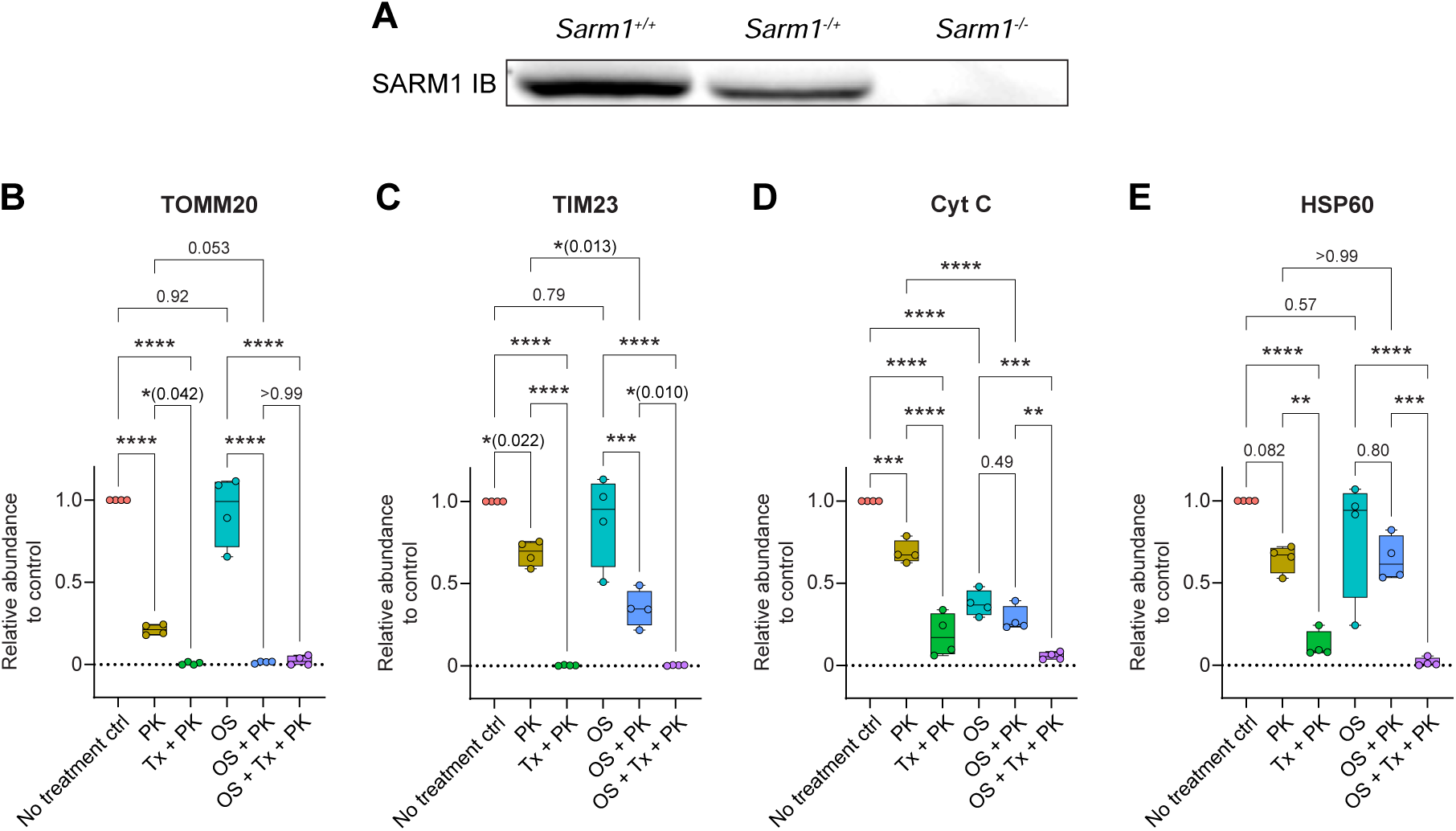
Western blot analysis of SARM1 and mitochondrial proteins (A) Validation of the SARM1 antibody in *Sarm1^+/+^*, *Sarm1^-/+^* and *Sarm1^-/-^* whole brain lysates. (B-E) Quantification of TOMM20, TIM23, Cyt C and HSP60 in the Proteinase K protection assay from the experiments shown in Figures 7C-D. N = 4 mice from 4 experiments. One-way ANOVA followed by Tukey’s multiple comparisons. Box plots denote minimum, first quartile, median, third quartile, and maximum values. \**P* < 0.05; \*\**P* < 0.01; \*\*\**P* < 0.001; \*\*\*\**P* < 0.0001.

## Notes

### Competing Interest Statement

The authors have declared no competing interest.

